# Programmed ribosomal frameshifting during *PLEKHM2* mRNA decoding generates a constitutively active proteoform that supports myocardial function

**DOI:** 10.1101/2024.08.30.610563

**Authors:** Gary Loughran, Raffaella De Pace, Ningyu Ding, Jianchao Zhang, Irwin Jungreis, Gionmattia Carancini, Jonathan M. Mudge, Ji Wang, Manolis Kellis, John F. Atkins, Pavel V. Baranov, Andrew E. Firth, Xiaowei Li, Juan S. Bonifacino, Yousuf A. Khan

**Affiliations:** School of Biochemistry and Cell Biology, University College Cork, Cork, Ireland; Neurosciences and Cellular and Structural Biology Division, Eunice Kennedy Shriver National Institute of Child Health and Human Development, National Institutes of Health, Bethesda, MD 20892, USA; Department of Cardiology, The First Affiliated Hospital of Zhengzhou University, Zhengzhou, Henan, Republic of China and Henan Key Laboratory of Hereditary Cardiovascular Diseases, Zhengzhou 450052, China; MIT Computer Science and Artificial Intelligence Laboratory, Cambridge, MA, USA; Broad Institute of MIT and Harvard, Cambridge, MA, USA; European Molecular Biology Laboratory, European Bioinformatics Institute, Wellcome Genome Campus, Hinxton CB10 1SD, Cambridge, UK; Department of Pathology, University of Cambridge, Cambridge, UK; Institute of Precision Medicine, The First Affiliated Hospital, Sun Yat-sen University, Guangzhou 510080, Republic of China; Department of Molecular and Cellular Physiology, Stanford University, Stanford, CA, USA; Department of Neurology and Neurological Sciences, Stanford University, Stanford, CA, USA; Department of Structural Biology, Stanford University, Stanford, CA, USA; Department of Photon Science, Stanford University, Stanford, CA, USA

## Abstract

Programmed ribosomal frameshifting is a process where a proportion of ribosomes change their reading frame on an mRNA^1^, rephasing the ribosome relative to the mRNA. While frameshifting is commonly employed by viruses^2^, very few phylogenetically conserved examples are known in nuclear encoded genes and some of the evidence is controversial^3,4^. Here we report a +1 frameshifting event during decoding of the human gene *PLEKHM2*^5^. This frameshifting occurs at the sequence UCC_UUU_CGG, which is conserved in vertebrates and is similar to an influenza virus sequence that frameshifts with similar efficiency^6,7^. The new C-terminal domain generated by this frameshift forms an α-helix, which relieves PLEKHM2 from autoinhibition and allows it to move to the tips of cells *via* association with kinesin-1 without requiring activation by ARL8. Reintroducing both the canonically-translated and frameshifted protein are necessary to restore normal contractile function of *PLEKHM2*-knockout cardiomyocytes, demonstrating the necessity of frameshifting for normal cardiac activity.

## Main

Ribosomes catalyze the synthesis of proteins, a tightly coordinated, regulated, and conserved process^8^. During elongation, each codon of an mRNA is matched to its amino acid-specific tRNA. The nascent peptide chain is then constructed in a sequential manner as tRNAs continue to deliver amino acids^9^. Maintenance of the reading frame is critical to ensure that proteins are correctly synthesized. A network of interactions between the ribosome, tRNAs, and other factors help to maintain translation fidelity^10^. Thus, the rate of spontaneous frameshifting, where the ribosome shifts reading frame during elongation, is exceptionally low^11^.

Programmed ribosomal frameshifting (PRF) occurs when a proportion of elongating ribosomes are induced to shift their reading frame by one or two nucleotides upon encountering specific signals in *cis-* or a *trans*-acting factor^1^. *Cis*-acting PRF signals may include a slippery site where tRNAs can re-pair in an alternative reading frame, a 3′ RNA-element, or a nascent peptide that can interact with the ribosome’s peptide exit tunnel. *Trans*-acting stimulators of PRF include proteins^12,13^ and small molecules such as polyamines^14^. Except where a stop codon is immediately encountered, PRF results in ribosomes synthesizing a distinct polypeptide from a sequence 3′ of the PRF site. Many viruses, including SARS-CoV-2^15^ and HIV-1^16^, require PRF for their gene expression. In vertebrates, there are few known examples of PRF required for the expression of nuclear genes. All vertebrate −1 PRF cases described to date are of viral origin^17,18^ as reported cases of non-retrovirus-derived −1 PRF in vertebrate genes were shown to be artefacts^3,4^. The genes for antizymes – negative regulators of cellular polyamine levels – require +1 PRF to synthesize full-length antizyme proteins, and the frameshifting provides an autoregulatory circuit where the efficiency of PRF increases in response to elevated polyamine levels^14^. While examples of translation from long overlapping reading frames in vertebrates are known^19,20^, these result from leaky scanning or alternative splicing rather than PRF. An influenza virus-like +1 PRF site has been bioinformatically predicted in the vertebrate gene *ASXL1*, although this has not been experimentally validated^21^. To date, there are no experimentally confirmed PRF signals in vertebrate cellular genes that provide ribosome access to internally overlapping ORFs.

Here, we report a highly conserved +1 PRF site in the mRNA encoding PLEKHM2 (also known as SKIP) that mediates ARL8-dependent coupling of lysosomes to the anterograde microtubule motor kinesin-1^22^. PRF results in the synthesis of a transframe protein, PLEKHM2-frameshift (FS). This protein does not contain the autoinhibitory C-terminal domain of the canonically translated protein product, PLEKHM2-CT^23^, and instead is predicted to contain an α-helical domain that we found to promote self-association. This results in a proteoform that does not require activation by ARL8 to promote kinesin-1-driven lysosome transport. We then infected *PLEKHM2*-knockout human induced pluripotent stem cells that were differentiated into cardiomyocytes (hiPSC-CMs) with constructs either encoding for the PLEKHM2-CT, PLEKHM2-FS, or both proteoforms. We found that reintroducing only the PLEKHM2-CT proteoform did not significantly recover contractility activity whereas reintroducing a combination of both the CT and FS proteoforms resulted in significant recovery.

### *PLEKHM2* expression involves a highly conserved +1 PRF

During bioinformatic scans for alternative ORFs, we noticed a long, conserved overlapping ORF in one of the coding exons of *PLEKHM2* (**Fig. 1a**). Our extensive surveying of transcriptomics datasets did not provide evidence that this overlapping frame can be accessed by translation of a short RNA isoform produced with alternative splicing or alternative transcriptional initiation. Towards the 5′ end of this alternative ORF is a potential +1 PRF sequence, UCC_UUU_CGG, almost identical to the influenza A virus UCC_UUU_CGU +1 PRF site, that is conserved in vertebrate *PLEKHM2* mRNAs (**Fig. 1b**). For the influenza A virus PRF, slippage is thought to occur when UUU is positioned in the ribosomal P-site and CGU is positioned in a presumably empty A-site^6^. Both UUU and UUC are decoded by the same tRNA isoacceptor whose anticodon, 3′-AAG-5′, has a higher affinity for UUC in the +1 frame than for the zero-frame UUU^24^. Subjecting *PLEKHM2* transcript sequence alignments to both Synplot2^24^ and MLOGD^26^ analysis, we found that, in mammals, sauropsids, amphibians and teleost fish, in the region of the *PLEKHM2* gene that is overlapped by the *FS* ORF, there is statistically significantly enhanced synonymous site conservation, and a positive coding signature specifically in the +1 reading frame (**Fig. 1c-f**).

**Figure 1.**
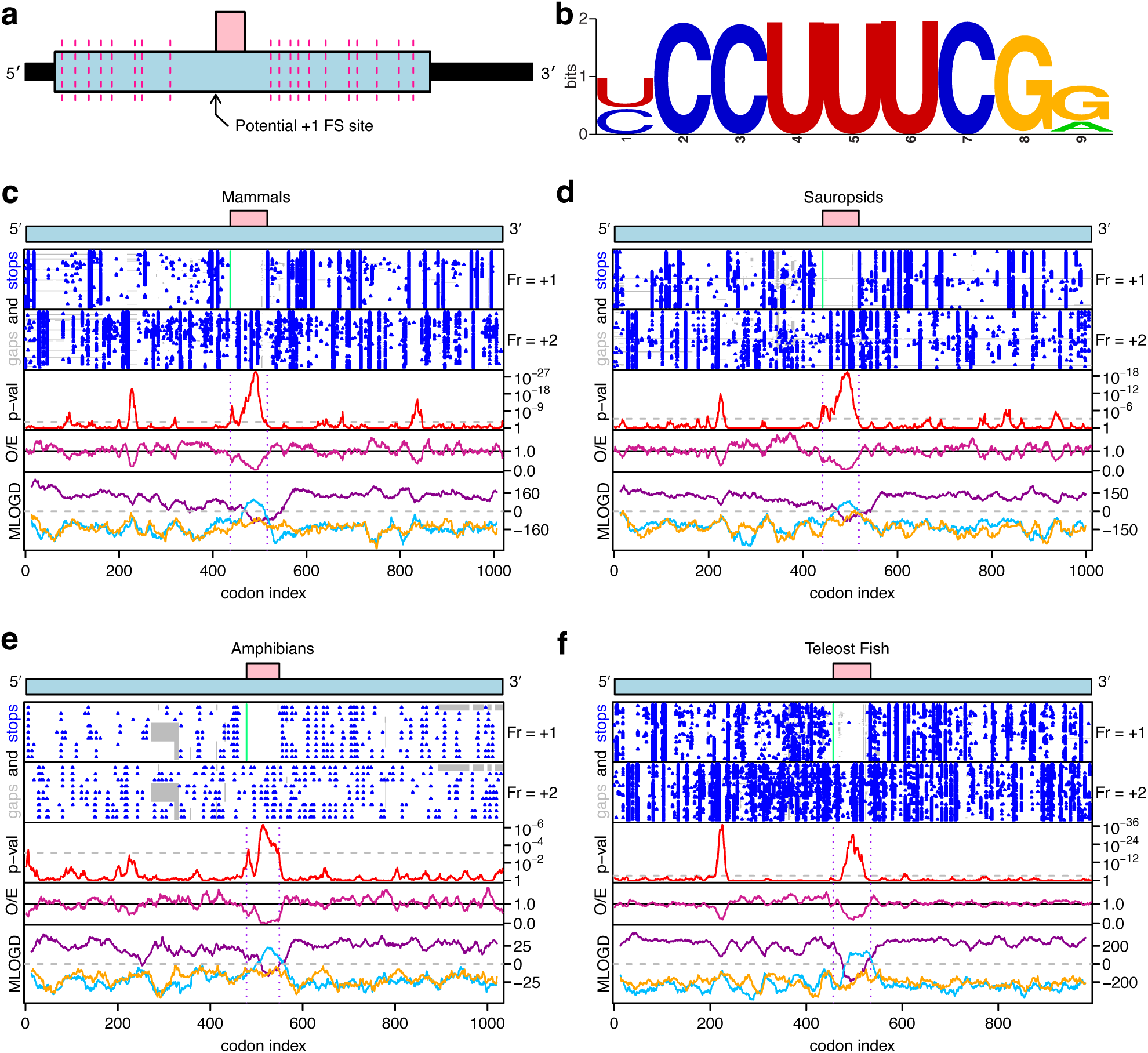
Synonymous site conservation and coding potential within the *PLEKHM2* dual coding region. **a.** Schematic of the main *PLEKHM2* transcript. Pink-dashed lines indicate exon-exon junctions and black bars represent untranslated regions (UTRs). The predicted PRF cassette is indicated with an arrow. The pink rectangle represents the +1 frame, conserved ORF. **b.** Sequence logo of putative PRF cassette, generated from alignment of *PLEKHM2* mRNA sequences from mammals, sauropsids, amphibians and teleost fish. **c-f.** Synplot2 and MLOGD plots for mammals (panel c), sauropsids (panel d), amphibians (panel e) and teleost fish (panel f). Top schematic shows stop codons (blue) in aligned sequences in +1 frame. Next row shows stop codons in +2 frame. O/E plot shows ratio of observed to expected synonymous substitutions under a null model of neutral evolution at synonymous sites and p-val plot shows corresponding p-value. Bottom row shows MLOGD coding potential plot for all three frames (purple 0, blue +1, yellow +2).

PhyloCSF, an algorithm used to predict conserved coding genomic regions, measures protein coding potential by computing the log-likelihood of a multispecies alignment under coding and noncoding models of evolution^27^. The PhyloCSF tracks in the UCSC Genome Browser indicate a conserved change of reading frame in exon 9 of *PLEKHM2* followed by coding resumption in the original reading frame downstream of the *FS* stop codon, in both mammals and birds (**Fig 2a,b**), suggesting a potential PRF^28^. A whole genome alignment of 100 vertebrate species rendered using CodAlignView (https://data.broadinstitute.org/compbio1/cav.php) shows that (i) both the CC_UUU_C 6-mer in the key portion of the potential PRF cassette and the *FS* stop codon are present in all aligned species, (ii) there is a preponderance of synonymous substitutions relative to the +1 frame in the latter portion of the potential dual coding region, indicating purifying selection on the amino-acid sequence translated from that frame, and (iii) there are no stop codons in the +1 frame within the potential dual coding region despite many stop codons in the flanking sequences on both sides of the aligned sequences (**S1a-d**).

**Figure 2.**
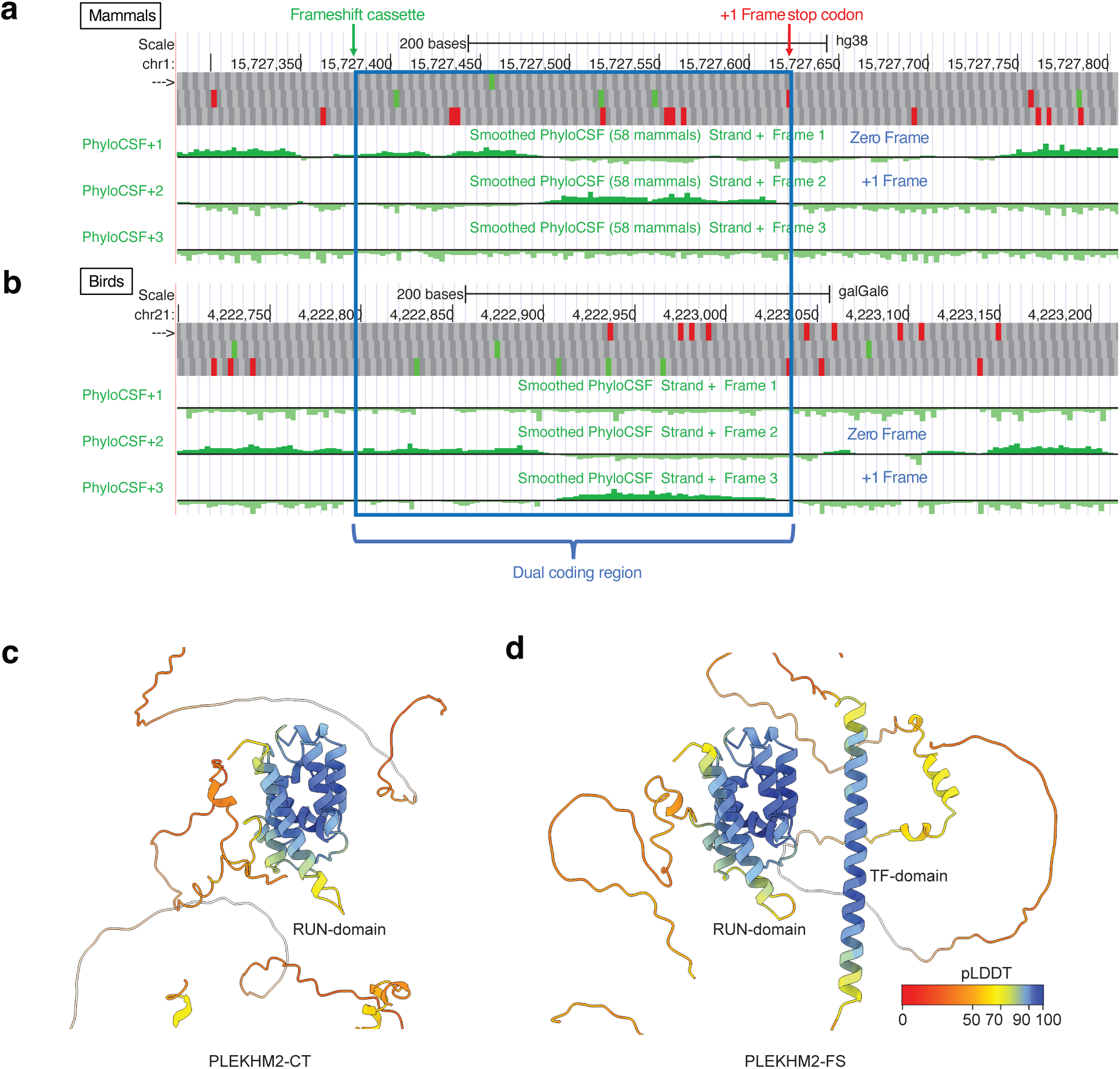
PhyloCSF and AlphaFold2 predictions on PLEKHM2. **a-b.** UCSC Genome Browser tracks showing evolutionary signature of protein coding potential in each of three reading frames as measured by PhyloCSF in mammals (**a**) and birds (**b**). Part way through the dual coding region (blue) between the PRF cassette (green arrow) and the +1 frame stop codon (red arrow), the PhyloCSF signal changes to the +1 frame, indicating greater amino acid conservation in the +1 frame, then resumes in the zero frame downstream of the +1 frame stop codon. **c.** AlphaFold2 predictions of PLEKHM2-FS using the standard workflow (left) and by generating MSAs with trans-frame sequence (right). pLDDT coloring scheme is shown in the bottom right.

Given that the protein sequence encoded by the overlapping ORF is highly conserved, we were curious if it encoded a protein with a high-confidence predicted structure. Using ColabFold’s^29^ implementation of AlphaFold2^30^, and assuming PRF on UCC_UUU_CGG, we input the sequence of the predicted FS protein. While ColabFold reliably predicted the N-terminal portion of the CT protein (**Fig. 2c**), it was unable to predict a structure from the portion of the protein encoded by the alternative ORF. However, given that AlphaFold2 relies on multiple sequence alignments (MSAs) from available databases to extract critical coevolutionary information, we reasoned that it would not have *a priori* knowledge on PLEKHM2-FS since its C-terminal sequence is absent from all existing databases. We generated custom MSAs for PLEKHM2-FS across all vertebrates, injected them into the model and were able to predict a high confidence α-helical domain in the transframe portion of the protein, further suggesting that this was a bona fide +1 PRF signal that enabled expression of a functional transframe protein (**Fig. 2d**).

### Experimental evidence of +1 PRF in *PLEKHM2*

Possible PRF into the +1 reading frame was tested experimentally by inserting the putative frameshift site and flanking sequence from *PLEKHM2* between *Renilla* and firefly luciferase CDS (coding sequence)s such that a +1 PRF event would give rise to a ∼100-kDa fusion product of the two reporter polypeptides^31,32^. These constructs were transfected into HEK293T cells and cell lysates were immunoblotted with anti-*Renilla* antibody. This demonstrated PRF in a WT construct, but not upon mutation of the putative PRF cassette from UCC_UUU_CGG to agC_UUc_aGa (**Fig. 3a**).

**Figure 3.**
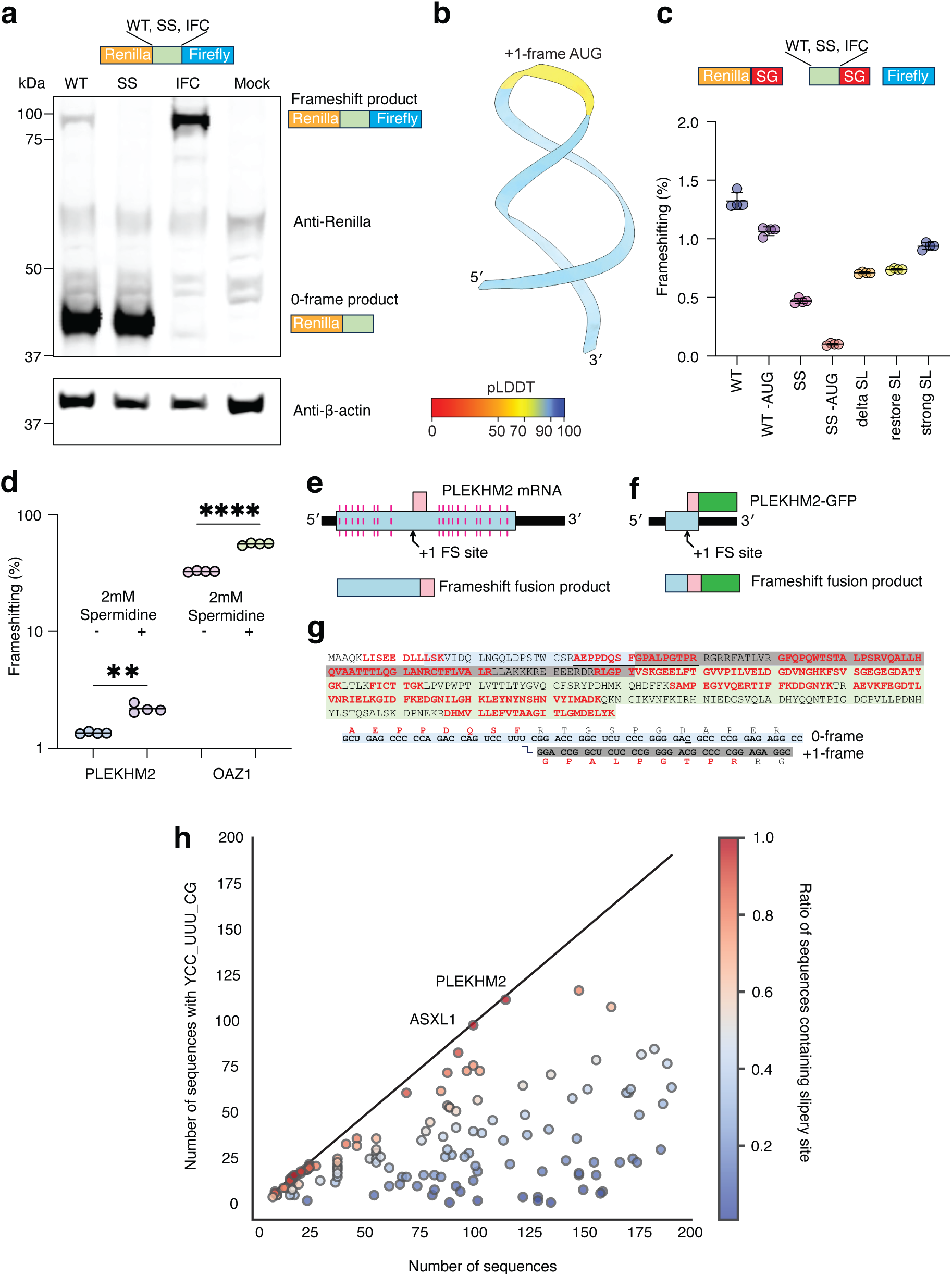
PRF on *PLEKHM2* mRNA. **a.** Representative immunoblot of whole cell lysates prepared from HEK293T cells transfected with *PLEKHM2* dual luciferase constructs. WT: wild-type (UCC_UUU_CGG), SS: frameshift mutant (agC_UUc_aGa), IFC: in-frame control (agC_UUc_Ga) with both reporters in the zero-reading frame, MOCK: no plasmid transfected. **b.** AlphaFold3 prediction of downstream RNA structure to the slippery site. RNA backbone is shown in a cartoon representation with nucleotides colored by pLDDT. The three nucleotides that show low pLDDT confidence (yellow) are the +1-frame AUG codon. **c.** PRF efficiencies (%) of *PLEKHM2* determined by dual luciferase assays. WT: wild-type (UCC_UUU _CGG), SS: frameshift mutant (agC_UUc_aGa). Change of +1 frame AUG to ACG labeled as ‘-AUG’. Delta SL, restore SL and strong SL changes are depicted in Supplemental Data Fig. 2b. *n* = 4. **d.** PRF efficiencies for *PLEKHM2* and *OAZ1* PRF cassettes upon addition of 2 mM spermidine (SPD). *n* = 4. **e.** Schematic depicting how mRNA of *PLEKHM2* corresponds to frameshifted fusion product. **f.** Schematic of mass-spectrometry construct. **g.** Mass-spectrometry results, with identified peptides highlighted in red font and junction peptide underlines. Junction peptide coverage shown below, with line depicting the transition from 0-to +1-frame, confirming the site of PRF. **h.** Plot depicting the degree to which human genes containing +1 PRF motif YCC_UUU_CG are conserved across sequence alignments. Points are colored by the fraction of aligned sequences of a gene containing the +1 PRF motif.

To measure the PRF efficiency of *PLEKHM2*, we used a dual luciferase reporter system that we previously described^33^. Briefly, the test sequence with its downstream sequence (**Fig. 3b**) is flanked by tandem StopGo sequences from foot-and-mouth disease virus^34^ that, while allowing continued protein synthesis, prevent peptide bond formation at a specific site in its sequence, resulting in expression of the reporter as separate proteins (**Fig. 3c**). This arrangement avoids potential artefacts that can arise when the test sequence fusion alters the individual reporter activities or stabilities^3,4,33^. Furthermore, we changed a +1 frame AUG within the *PLEKHM2* cassette to minimize the possibility of firefly luciferase translation due to initiation on potential low abundance mRNAs that may be generated by cryptic splicing or cryptic promoters. Dual luciferase assays on lysates prepared from HEK293T cells transfected with *PLEKHM2* constructs indicated a PRF efficiency of ∼1.3% for WT and ∼1.1% for WT minus AUG (**Fig. 3c** and **Ext 1a,b**). The slight decrease in firefly luciferase activity upon mutation of the +1 frame AUG indicates some level of internal initiation on this codon, underscoring the importance of controlling for such artefacts.

Although +1 PRF by mammalian ribosomes on influenza virus-like shift sites, such as in influenza A virus and *PLEKHM2*, do not appear to require *cis*-acting stimulators, we noticed a stem-loop 3′ of the *PLEKHM2* shift site that appears to display sequence conservation (**Fig. S2a**) and was predicted confidently by AlphaFold3^35^ (**Fig. 3b**). To test the possible stimulatory role of this putative stem-loop structure, we introduced mutations to disrupt (delta SL), restore (restore SL) or strengthen (strong SL) the stem-duplex (**Fig. S2b**). Disrupting the putative stem-loop structure decreased *PLEKHM2* PRF efficiency to ∼0.75%. However, WT levels of PRF were not restored by changes that were predicted to restore stem-loop formation or strengthen the stem-loop (**Fig. 3c** and **Ext 1a,b**). Although +1 PRF on the *PLEKHM2* mRNA does not appear to require the predicted stem-loop RNA secondary structure, there may be a weak stimulatory effect. It is possible that in certain cell types or under certain conditions the RNA secondary structure may modulate +1 PRF in *PLEKHM2* expression.

Polyamines stimulate the translation of the full antizyme 1 protein (encoded by *OAZ1*) by using +1 PRF to bypass an early stop codon in the zero-frame^36^, a mechanism conserved from protists to mammals. Therefore, we tested whether +1 PRF in the *PLEKHM2* gene may also be stimulated by polyamines, since polyamines have been proposed to stimulate +1 PRF in a sequence-independent manner^37,38^. HEK293T cells transfected with *PLKEHM2* or *OAZ1* dual luciferase constructs were treated with spermidine and assayed for PRF efficiency. Although baseline *OAZ1* PRF (32.5%) is much higher than *PLEKHM2* (∼1 %), we observed a similar 1.5-fold increase in PRF efficiency for both *OAZ1* and *PLEKHM2* when spermidine was added (**Fig. 3d** and **Ext. 1c,d**). This indicates that *PLEKHM2* PRF efficiency can be regulated. To assess whether *PLEKHM2* PRF may be more efficient in other cells or tissues, we compared ribosome read densities extracted from publicly available ribosome profiling data, upstream and downstream of the PRF casette in the *PLEKHM2* mRNA. Although subtle differences in read densities were observed among different tissues, the baseline rate of PRF is too low to determine if these differences are significant (**Fig. S3**).

To determine the precise site and direction of PRF, we aimed to identify transframe peptides by mass spectrometry. We generated a construct in which a CDS encoding green fluorescent protein (GFP) was fused in-frame to the 3′ end of the transframe CDS (**Fig. 3e,f**). An additional construct (PLEKHM2-K-GFP) introducing a lysine codon close to the slip site was used to increase the chances of identifying the junction tryptic peptide (**Ext. 1e**). Synonymous mutations in the zero-frame were introduced to alter two +1 frame AUG codons to avoid potential initiation in the +1 frame. Frameshift expression from these constructs would result in the transframe fusion PLEKHM2-GFP or PLEKHM2-K-GFP, which could be affinity-purified on GFP-Trap beads, while non-frameshift expression would result in products that do not contain GFP. Constructs were expressed in HEK293T cells, and transframe fusions were affinity-purified from cell lysates and resolved by SDS-PAGE. Both constructs directed synthesis of a specific protein migrating at the predicted molecular weight for PLEKHM2-FS-GFP (∼34 kDa). Gel slices containing these proteins were excised, digested with trypsin, and the resulting peptides were analyzed by liquid chromatography tandem mass spectrometry (LC-MS/MS). Tryptic peptides covering 59% of the PLEKHM2-FS-GFP construct and 85% of the PLEKHM2-K-FS-GFP construct were identified, including peptides encoded both upstream and downstream of the shift site (**Fig. 3g and Ext. 1e**). Importantly, three peptides spanning the shift site itself were identified. These peptides, AEPPDQSFGPALPGTPR (from PLEKHM2-FS-GFP), QSFGPALPGTPR and AEPPKQSFGPALPGTPR with a missed tryptic cleavage (from PLEKHM2-K-FS-GFP), define the shift site (UCC_UUU_CGG) and direction (+1) of PRF.

Given that the YCC_UUU_CG +1 PRF motif clearly stimulates significant levels of frameshifting in *PLEKHM2* (**Fig. 3c**) and influenza A virus^6,7^ without the necessity of a downstream RNA structure, we then investigated the presence of this motif in other human genes. Starting with 17,922 sequence alignments of human CDSs with their mammalian (or sometimes other chordate) homologues selected at a ≥65% amino acid identity threshold (**Methods**), we found 134 occurrences of YCC_UUU_CG in human CDSs. The degree and phylogenetic extent of conservation of this motif in the *PLEKHM2* sequence alignments is exceptional (**Fig. 3h**). Nonetheless, there are multiple other occurrences of YCC_UUU_CG in human CDSs (such as ASXL1, which also shows exceptional conservation), and these would also be expected to give rise to low level frameshifting. Given this, there is potential that phylogenetically restricted (e.g. primate-specific or even human-specific) frameshifting at these other sites occurs.

### The PLEKHM2-FS proteoform is constitutively active

Next, we investigated the activity of the PLEKHM2-FS proteoform. PLEKHM2-CT^5^ comprises a RUN domain that interacts with the lysosome-associated small GTPase ARL8^22^, WD-WE motifs that interact with kinesin-1^39^, a largely disordered region, and three autoinhibitory PH domains^23^ (**Fig. 4a**). ARL8 both recruits PLEKHM2-CT to lysosomes^22^ and relieves PLEKHM2-CT autoinhibition^23^. These properties enable PLEKHM2-CT to function as a regulated adaptor for ARL8-dependent coupling of lysosomes to kinesin-1 and consequent transport toward the plus end of microtubules^23,39^.

**Figure 4.**
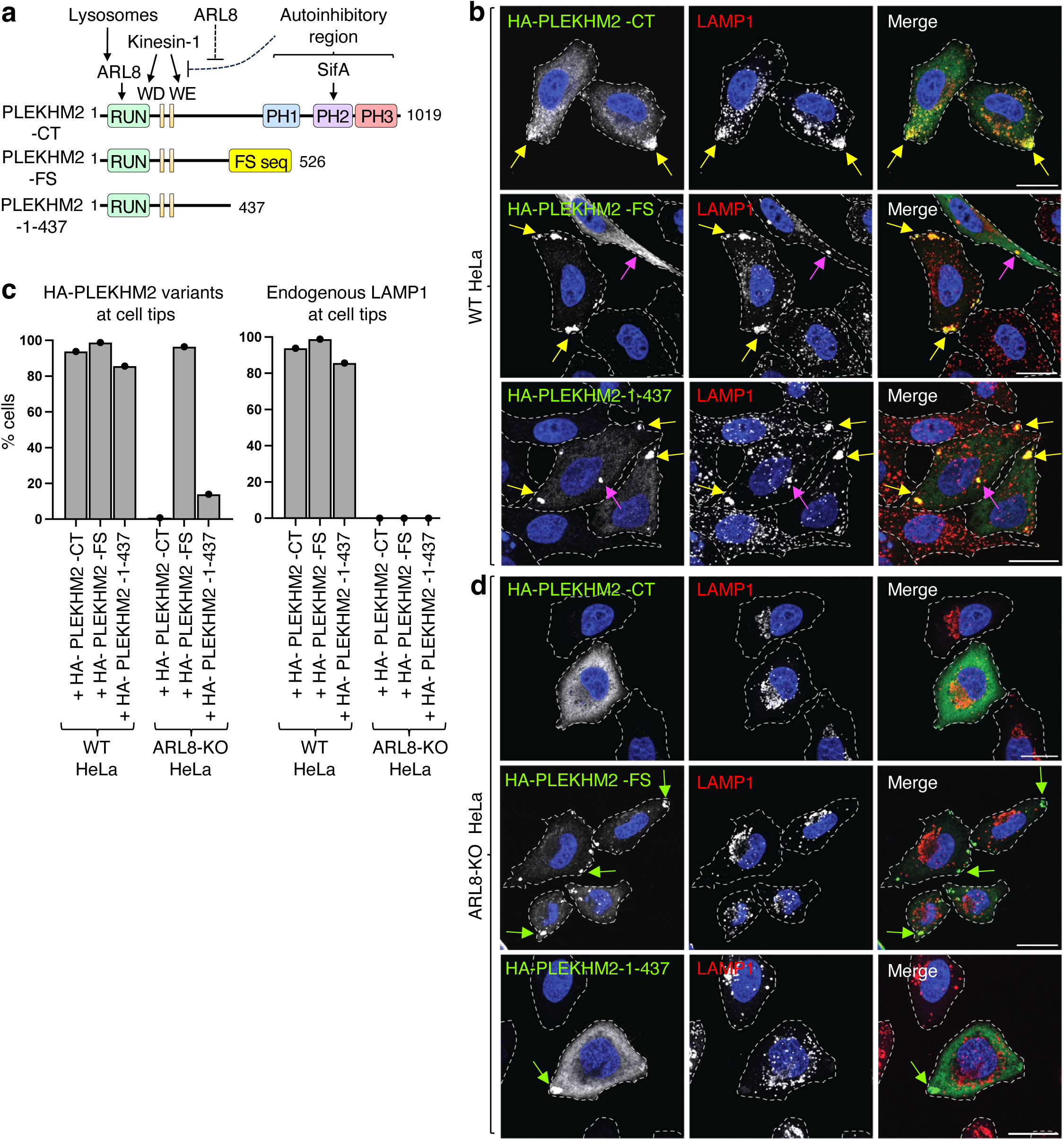
The PLEKHM2-FS proteoform is constitutively active in driving lysosome movements towards the cell periphery. **a.** Schematic representation of PLEKHM2 variants. The different domains/motifs are indicated by colored boxes: RPIP8, UNC-14, and NESCA (RUN); Trp-Asp (WD); Trp-Glu (WE); pleckstrin homology (PH). Arrows and dashed lines indicate positive and negative interactions, respectively, with binding partners. Amino-acid numbers are also indicated. **b-d**. WT HeLa cells were transiently transfected with the indicated HA-tagged PLEKHM2 constructs for 48 h and analyzed by confocal immunofluorescence microscopy for the HA epitope (green) and endogenous LAMP1 (lysosomes, red). Nuclei were labeled with DAPI (blue). Cell edges were outlined by staining of actin with Alexa Fluor™ 647-conjugated phalloidin (not shown) and indicated by dashed lines. Arrows in panels **b** and **d** indicate clusters of PLEKHM2 together with lysosomes (yellow arrows) or without lysosomes (green arrows) at cell tips or near the cell center (magenta arrows). Scale bars: 20 μm. Panel **c shows a** quantification of the percentage of WT (panel **b**) and ARL8-KO cells (panel **d**) displaying accumulation of PLEKHM2 variants or LAMP1 at cell tips. More than 150 cells per sample were scored. **d.** ARL8-KO HeLa cells were transfected and analyzed as described for panel **b**. Scale bars: 20 μm. **e.** Schematic representation of the BORC-ARL8-dependent activation and recruitment of PLEKHM2 to a lysosome (top), and the recruitment of the constitutive, ARL8-independent activity but ARL8-dependent recruitment of PLEKHM2-FS to a lysosome (bottom).

The PRF that generates PLEKHM2-FS preserves the RUN domain and WD-WE motifs but parts of the disordered region and the three PH domains are absent (**Fig. 4a**). As previously shown^23,39^, overexpression of HA-tagged PLEKHM2-CT (HA-PLEKHM2-CT) in WT HeLa cells (**Ext. 2a**) results in the redistribution of a large fraction of lysosomes immunolabeled for the lysosomal markers LAMP1 (**Fig. 4b,4c** and **Ext. 3a**) and LAMTOR4 (**Ext. 2b**) to cell tips (also referred to as vertices; indicated by yellow arrows). Overexpression of HA-PLEKHM2-FS or a truncated HA-PLEKHM2 1-437 lacking the three PH domains and the FS sequence (**Fig. 4a**, **Ext. 2a**) similarly redistributed lysosomes to cell tips (**Fig. 4b, 4c**, **Ext. 2b, 3a**). All these HA-PLEKHM2 variants themselves redistributed to cell tips along with lysosomes. PLEKHM2– lysosome clusters exhibited a tighter appearance and were occasionally observed to be more centrally located for the HA-PLEKHM2-FS and HA-PLEKHM2-1-437 constructs compared to the HA-PLEKHM2-CT construct (**Fig. 4b**, **Ext. 2b, 3a**; centrally located clusters are indicated by magenta arrows), a phenotype that likely stems from the removal of the C-terminal sequences. These observations thus demonstrated that PLEKHM2-FS is at least as active as PLEKHM2-CT, and that this activity does not depend on the FS sequence, when overexpressed in WT cells.

Knock out (KO) of both the ARL8A and ARL8B paralogs in HeLa cells (referred to as ARL8 KO) prevented the association of HA-PLEKHM2-CT with lysosomes (due to loss of ARL8-dependent recruitment) and kinesin-1 (due to loss of ARL8-dependent relief from autoinhibition), thereby impeding the overexpression-induced redistribution of both lysosomes and PLEKHM2 to cell tips (**Fig. 4c, 4d**, **Ext. 2a**). Importantly, although overexpressed HA-PLEKHM2-FS failed to redistribute lysosomes, it could itself relocate to the tips of ARL8-KO cells (**Fig. 4c, 4d**, **Ext. 2c**, green arrows). This latter phenotype differs from that observed for HA-PLEKHM2-CT and is likely attributable to the absence of the autoinhibitory PH domains in PLEKHM2-FS. These observations indicated that PLEKHM2-FS is constitutively active, requiring no activation by ARL8 for association with kinesin-1 and movement toward cell tips.

In ARL8-KO cells, the truncated HA-PLEKHM2-1-437 exhibited a more cytosolic distribution (**Fig. 4d**, **Ext 2c, S4a, S4b**) and substantially reduced movement to cell tips compared to PLEKHM2-FS (**Fig. 4c, 4d**, **Ext. 2c, 3b, S4a, S4b**). These observations suggest that the FS sequence does not just replace an autoinhibitory sequence but also enhances the interaction with kinesin-1. Co-immunoprecipitation of HA-and myc-tagged constructs showed that PLEKHM2-FS interacted with itself much more than with PLEKHM2-CT or PLEKHM2-1-437 (**Ext. 2d**). These data demonstrate that the constitutively active PLEKHM2-FS proteoform likely plays a role in remodeling the lysosome distribution in the cell via its enhanced interaction with kinesin-1, which is potentially mediated by the FS sequence’s propensity to self-associate.

### The PLEKHM2-FS proteoform contributes to cardiomyocyte contraction activity

Proper lysosomal function and positioning is critical for autophagy, a conserved process for bulk degradation and recycling of cellular components^40^. Autophagy plays an essential role in the heart^41^. Under normal conditions it is important for the turnover of proteins and organelles, such as in cardiomyocytes rich in mitochondria, which, when damaged, can release pro-apoptotic factors^42^. Previous studies have also shown that autophagy protects against ischemia/reperfusion injury in cardiac cells^43^ and is an adaptive response to protect cardiac cells from hemodynamic stress^44^. A two-base deletion mutation in PLEKHM2, a protein critical to lysosomal positioning and thus autophagy, has been identified in patients suffering from dilated cardiomyopathy (DCM) and left ventricular noncompaction (LVN)^45^. In fibroblasts derived from these patients, normal lysosomal distribution was perturbed, leading to impaired autophagy flux and thereby providing a mechanism for PLEKHM2’s role in myocardial contraction. Furthermore, disruption of PLEKHM2 splicing has also been shown to lead to DCM and subsequent death^46^. This led us to investigate the potential role of the PLEKHM2 proteoforms in cardiac function.

Given that the chambered heart system arose evolutionarily in vertebrates^47^, we wanted to see how conserved the PLEKHM2-FS protein sequence was relative to other key vertebrate proteins. By determining the set of all conserved proteins between *Homo sapiens* and *Danio rerio*, we were then able to pairwise compare every single protein sequence and determine a conservation distribution (**Fig. 5a, 5b**). PLEKHM2-FS, also conserved in all vertebrates, was more conserved than most key conserved vertebrate proteins (**Fig. 5b**), suggestive of its essential role in vertebrate biology.

**Figure 5.**
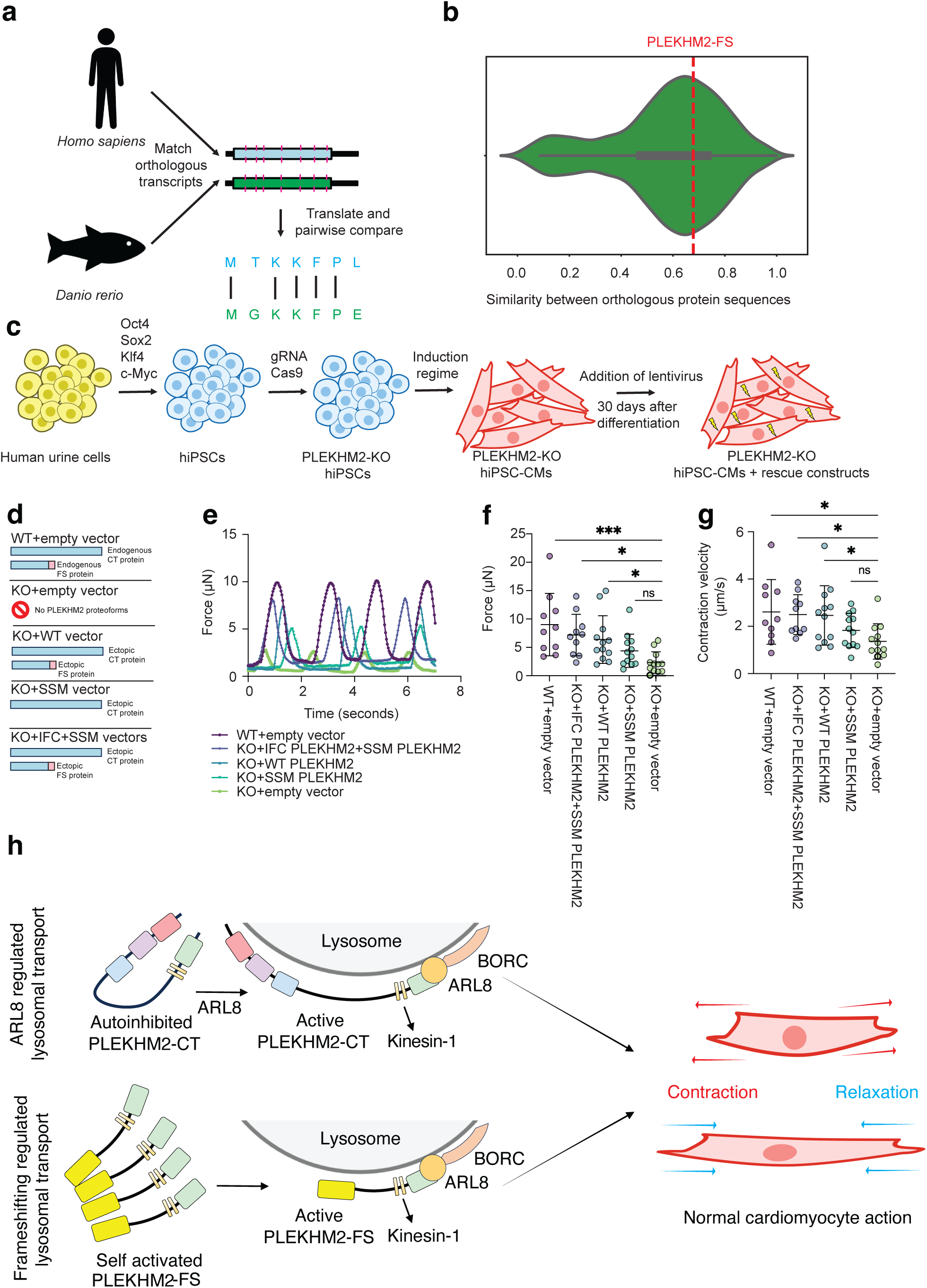
The PLEKHM2-FS proteoform contributes to cardiac function. **a.** Schematic representation of the generation of a set of conserved proteins between *H. sapiens* and *D. rerio* and computing pairwise orthology. **b.** Distribution of the similarity of conserved protein sequences in vertebrates, with a red line representing the degree of PLEKHM2-FS proteoform conservation. **c.** Schematic representation of the process of establishing hiPSC-CMs. **d.** Schematic representation of the different constructs used to infect hiPSC-CMs, illustrating which proteoforms (CT, FS) are expressed in the cell by each construct combination. **e.** Example of a single force trace experiment. **f-g.** Quantification of force and contraction velocity in WT hiPSC-CMs and PLEKHM2-KO hiPSC-CMs expressing different constructs analyzed with a one-way ANOVA (ns: p-value > 0.05, *: p-value < 0.05, **: p-value <0.01, ***: p-value <0.001). Each dot represents a set of measurements from an individual cell (n = 10-12). **h.** Schematic summarizing how PLEKHM2-CT and PLEKHM2-FS proteforms support cardiomyocyte function.

To further investigate PLEKHM2-FS’s physiological role, we used CRISPR generated PLEKHM2-KO human induced pluripotent stem cells (hiPSCs)^48^ (**Fig. 5c**, **methods**). These cells were then differentiated into cardiomyocytes (hiPSC-CMs)^49^. These hiPSCs-CMs were next infected with lentivirus containing either an empty vector, the WT PLEKHM2 cDNA, PLEKHM2 cDNA with a silent slippery site mutant (SSM), or with two vectors: SSM PLEKHM2 and PLEKHM2 cDNA with an in-frame control mutation (IFC). A control cell line where PLEKHM2 was not knocked out was also established and infected with an empty vector (**Fig. 5d**). As expected, the PLEKHM2-KO hiPSC-CMs showed ablated myocardial contractility (**Fig. 5e-g**, **S5a, S5b**), elevated levels of heart failure biomarkers NPPA and NPPB^50^ (**S5b, S5c**), abnormal calcium handling (**S5d-S5h**), and an accumulation of autophagy receptor p62/SQSTM1 (**S5i, S5j**), while most conditions reintroducing PLEKHM2 proteoforms to these PLEKHM2-KO hiPSC-CMs showed various degrees of recovery at or towards WT hiPSC-CMs.

Notably, however, PLEKHM2-KO hiPSC-CMs reinfected with the PLEKHM2 SSM vector, diminishing its ability to productively frameshift and therefore only producing the PLEKHM2-CT proteoform (**Fig. 3b**), were not able to significantly recover any aspect of myocardial contraction including force (**Fig. 5e, 5f**), contraction velocity (**Fig. 5g**), average contraction displacement (**S5a**) or relaxation velocity (**S5b**). Upon reintroduction of the PLEKHM2-CT proteoform along with the -FS proteoform, significant myocardial contraction was indeed restored (**Fig. 5e-g**, **S5a, S5b**), demonstrating the critical role that both the PLEKHM2-CT and PLEKHM2-FS proteoforms play in contraction.

Based on these data, we conclude that PLEKHM2-FS is a constitutively active variant of PLEKHM2 that does not require ARL8-dependent activation for kinesin-1–dependent movement of lysosomes towards the cell periphery. This proteoform, generated by +1 PRF with a slippery site similar to the one identified in influenza, is necessary along with the PLEKHM2-CT proteoform to promote myocardial contraction (**Fig. 5h**).

Although the conservation of PLEKHM2-FS in vertebrates coincides with the development of the modern, chambered heart, we do not rule out that PLEKHM2-FS may be involved in other physiological processes dependent on lysosomal distribution and therefore autophagy in the nervous system^51^, immunity^52^, and cancer^53^.

How the presence of this constitutively active form of PLEKHM2 specifically affects the dynamics of kinesin-1-driven lysosomal distribution within cells remains an open question. PLEKHM2-FS could provide a basal level of lysosome transport, while PLEKHM2 would enable another level of regulated lysosome transport in response to stimuli such as starvation^54^ or growth factor receptor activation^55^. Furthermore, PLEKHM2-FS may not just be constitutively activated, but could also play a role in activating the non-frameshift proteoform.

The ability of polyamines to similarly stimulate both *OAZ1* and *PLEKHM2* +1 PRF suggests substantial latitude in sequences at which polyamines stimulate +1 PRF. While polyamine stimulation of *OAZ1* PRF has evolved to maintain polyamine levels within a narrow cellular concentration, it is possible that polyamine stimulation of *PLEKHM2* PRF could be incidental – perhaps polyamines directly or indirectly affect elongation rates that is especially seen in PRF events.

The discovery of the first PRF signal in vertebrate cellular genes that provides ribosome access to internally overlapping ORFs thus marks a major milestone moving this phenomena beyond viruses and sets the stage for further searches in vertebrate genomes. Other conserved PRF signals may exist but may not have been as highly selected for or evolved as early as the one in *PLEKHM2*, making them less readily identifiable through traditional bioinformatic searches. Of the 134 additionally identified genes containing the verified +1 PRF motif, it is possible that a subset of them productively employ +1 PRF to encode an alternative protein product. Furthermore, as more PRF motifs are identified, more PRF employing genes will likely be identified in vertebrate genomes. The increasing number of viral and cellular examples of PRF signals will lend itself to complementary computational techniques, such as machine learning, and improved high-throughput assays.

**Extended Figure 1.**
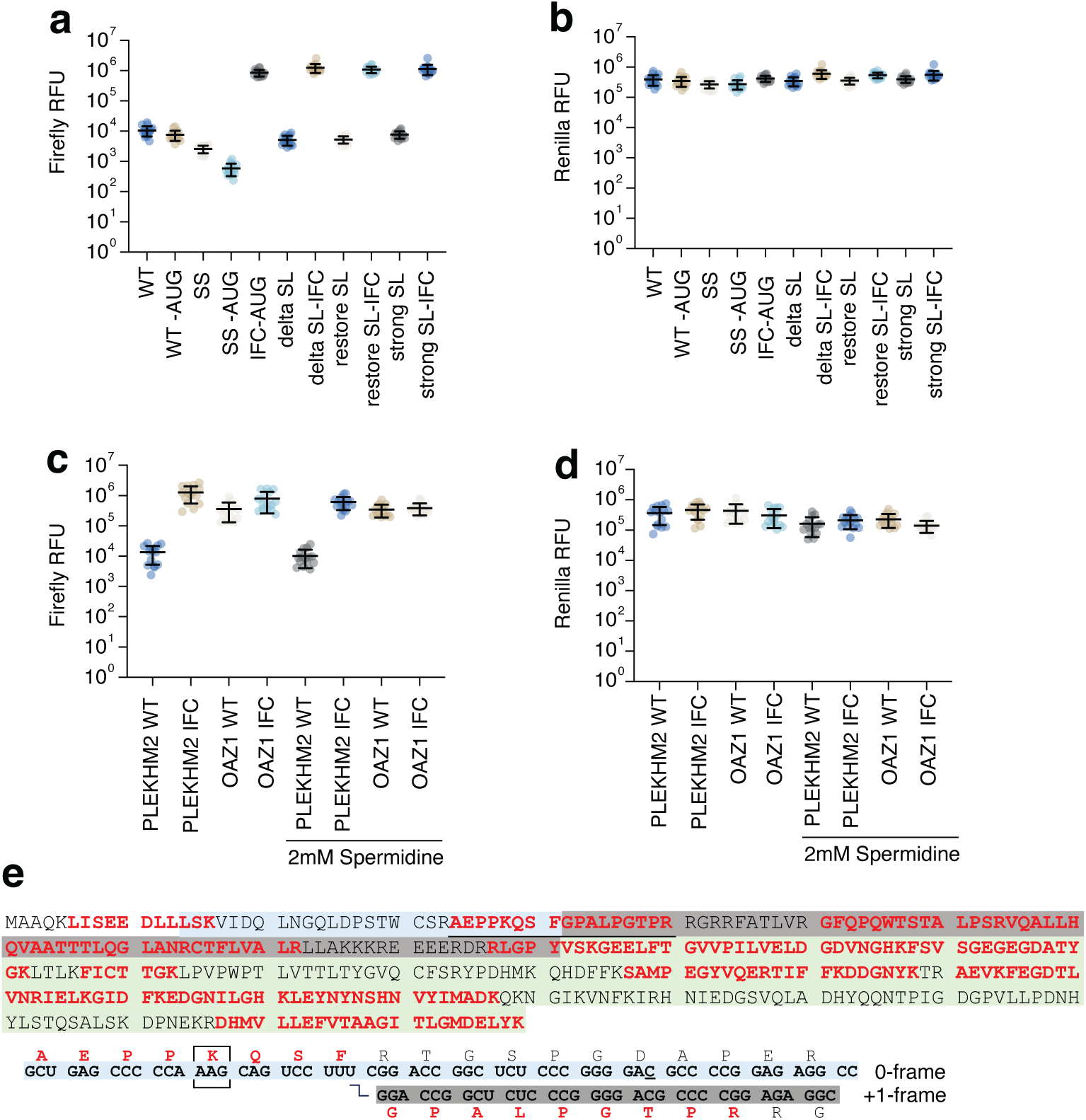
PLEKHM2 luciferase assays and additional mass-spectrometry data. **a.** Absolute firefly luciferase values. **b.** Absolute *Renilla* luciferase values; all values are within the same order of magnitude. **c.** Absolute firefly luciferase values in spermidine experiments. **d.** Absolute *Renilla* luciferase values in spermidine experiments. **e.** Mass-spectrometry results of PLEKHM2-K-GFP constructs; covered peptides are shown in red font with the junction peptide underlined. The transition of the reading frame is shown below.

**Extended Figure 2.**
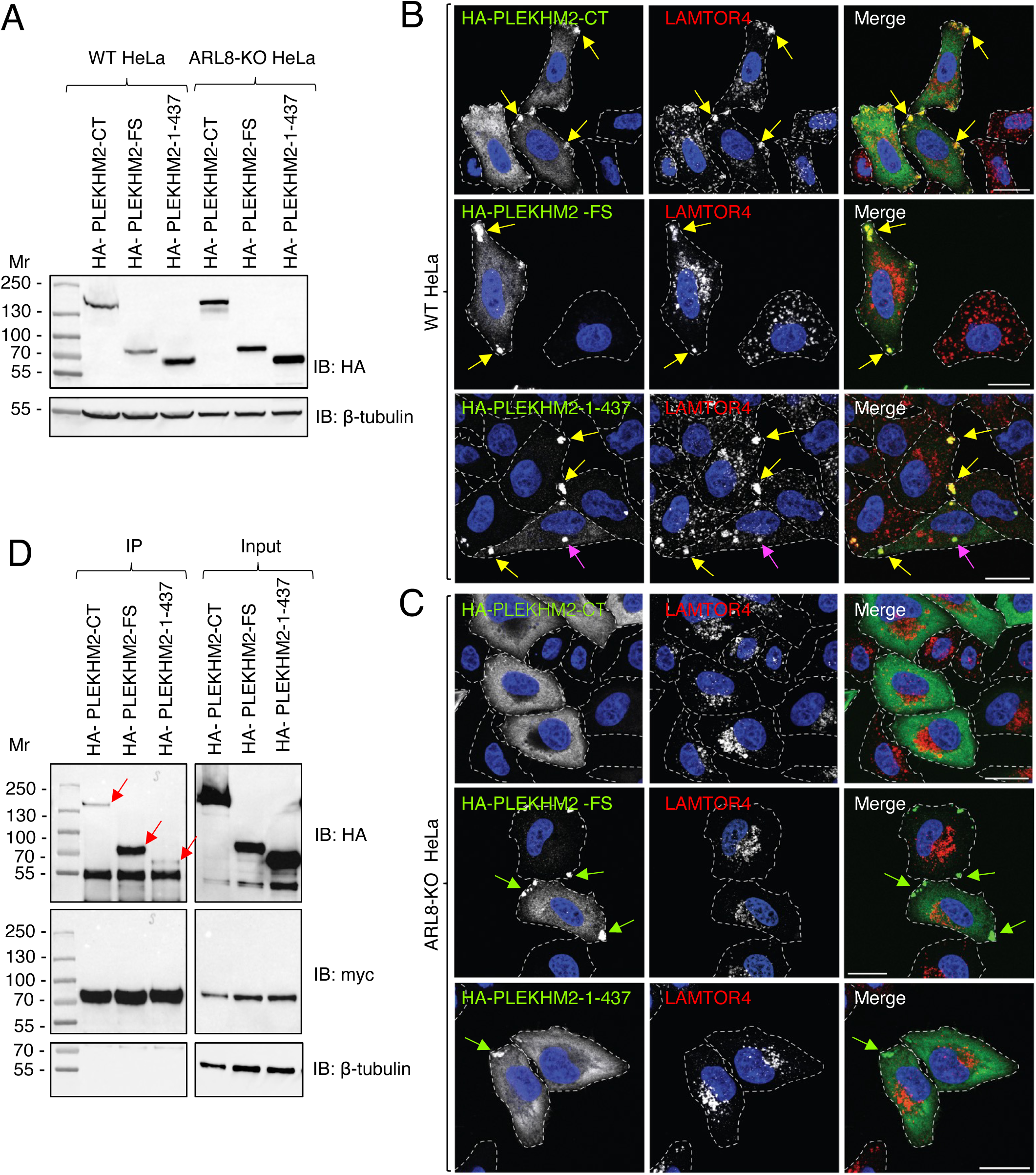
Expression, activity, and self-association of PLEKHM2-FS. **a.** WT or ARL8-KO HeLa cells were transiently transfected with the indicated HA-tagged PLEKHM2 constructs (see scheme in Fig. 4a) for 48 h and analyzed by SDS-PAGE and immunoblot analysis with antibodies to the HA epitope. β-tubulin was used as a loading control. The positions of molecular mass markers (Mr, in kDa) are indicated on the left. **b.** WT HeLa cells like those shown in panel **a** were analyzed by confocal immunofluorescence microscopy for the HA epitope (green) and endogenous LAMTOR4 (lysosomes, red). Nuclei were labeled with DAPI (blue). Cell edges were outlined by staining of actin with fluorescent phalloidin (not shown) and indicated by dashed lines. Arrows indicate clusters of HA-PLEKHM2 together with lysosomes (yellow arrows) or without (green arrows) at the cell tips or more centrally (magenta arrows). Scale bars: 20 μm. Notice how all HA-PLEKHM2 constructs drive distribution of both HA-PLEKHM2 and lysosomes to cell tips. **c.** ARL8-KO HeLa cells were transfected and analyzed as described for panel **b**. Scale bars: 20 μm. Notice that HA-PLEKHM2-CT is unable to localize to and drive lysosomes to cell tips in ARL8-KO cells. In contrast, HA-PLEKHM2-FS can localize to, but cannot drive lysosomes to cell tips. HA-PLEKHM2-1-437 is incapable of moving either protein to cell tips. **d.** HEK293T cells were co-transfected with plasmids encoding the indicated HA-tagged constructs with myc-PLEKHM2-FS and subjected to immunoprecipitation with an antibody to the myc epitope. Immunoprecipitates (IP) and cell extracts (10%, Input) were analyzed by SDS-PAGE and immunoblotting (IB) for the HA and myc epitopes, or β-tubulin (loading control). Red arrows indicate the HA constructs that are co-immunoprecipitated with the myc-PLEKHM2-FS construct. The positions of molecular mass markers (Mr, in kDa) are indicated on the left. Notice the much greater co-immunoprecipitation of myc-PLEKHM2-FS with HA-PLEKHM2-FS relative to HA-PLEKHM2-CT and HA-PLEKHM2-1-437.

**Extended Figure 3.**
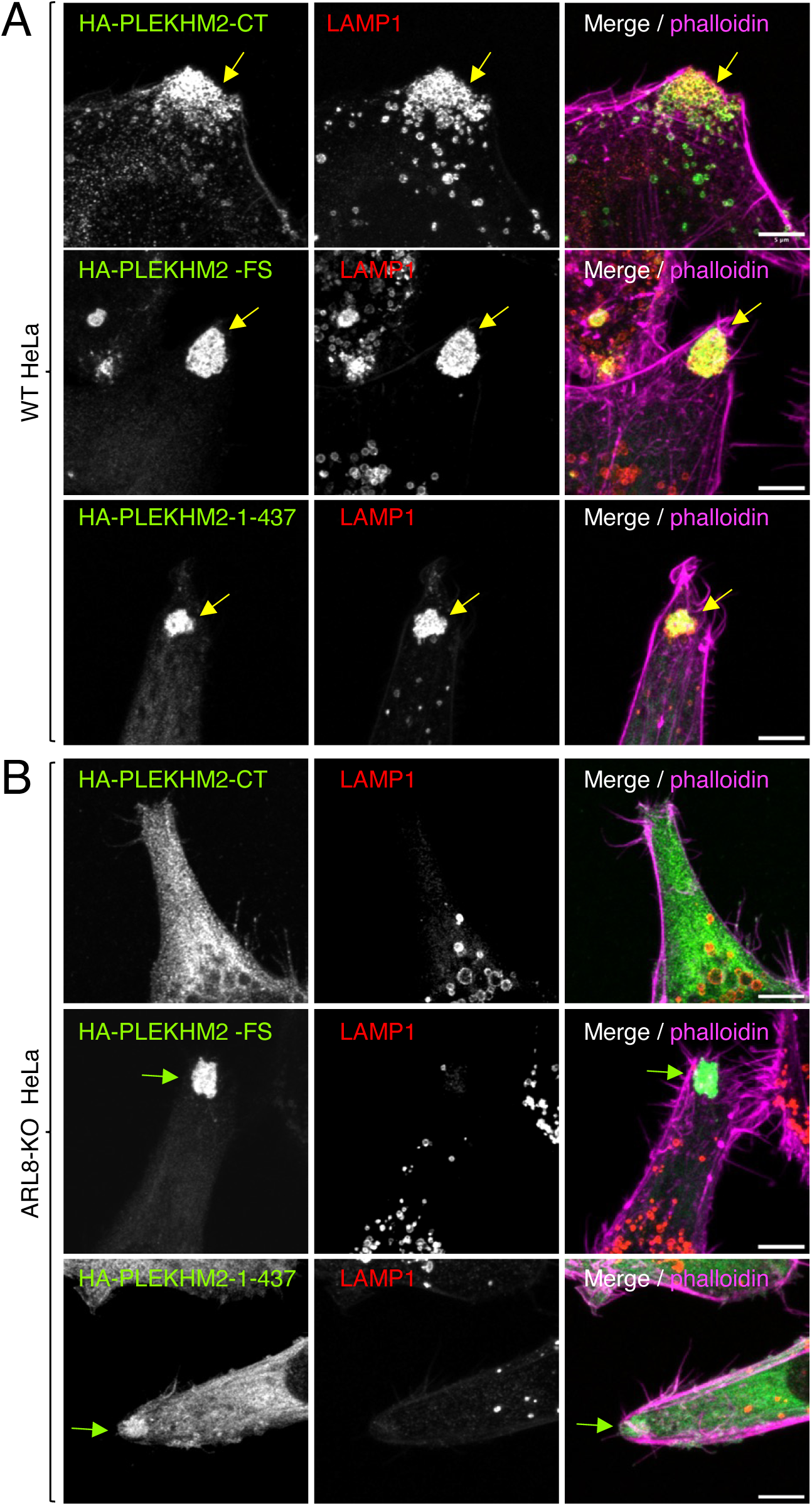
Airyscan microscopy of WT and ARL8-KO HeLa cells expressing different PLEKHM2 variants. **a.** WT HeLa cells were transiently transfected with the indicated HA-tagged PLEKHM2 constructs for 48 h and analyzed by immunofluorescence microscopy for the HA epitope (green), endogenous LAMP1 (red) and Alexa Fluor™ 647-conjugated phalloidin (magenta). Cells were imaged using a higher-resolution Airyscan confocal microscope. Arrows indicate clusters of PLEKHM2 together with lysosomes (yellow arrows) or without (green arrows) at cell tips outlined by Alexa Fluor™ 647-conjugated phalloidin. Scale bars: 5 μm. Notice that HA-PLEKHM2-FS or HA-PLEKHM2-1-437 form tight clusters with lysosomes. **b.** ARL8-KO HeLa cells were transfected, stained, and imaged as described for panel **a**. Notice the absence of HA-PLEKHM2 and the reduction of HA-PLEKHM2-1-437 at cell tips. Scale bars: 5 μm.

**Figure S1 | Genomic alignment of 100 vertebrate species in a neighborhood of the dual coding region, with features relevant to protein-coding evolution color-coded by CodAlignView.** For clarity, insertions relative to the human sequence are not shown in (a) and (b); the same alignments including these insertions are shown in (c) and (d), respectively. (a) Codonization and color coding with respect to the +1 frame. Preponderance of synonymous substitutions (light green) in the latter portion of the dual coding region indicates purifying selection on the amino acid sequence translated from this frame. There are many TGA (yellow), TAG (magenta), and TAA (cyan) stop codons in other species in this reading frame both 5’ of the PRF cassette (green arrow) and 3’ of the +1 frame stop codon (red arrow), but none within the dual coding region. The CCTTTC in the PRF cassette is perfectly conserved in all aligned species. (b) Same region codonized and color-coded with respect to the zero frame.

**Figure S2 | Potential RNA secondary structure downstream of slippery site**. **a.** Predicted RNA secondary structure using forna^56^ and CodAlignView with secondary structure regions highlighted. Uppercase letters in the diagram indicate which nucleotides were inputted for structure prediction. **b.** Diagram showing what structural mutants were made.

**Figure S3 | Ribosome profiling analysis mined from across different tissues**. Individual datasets are represented by dots coloured by the tissue of origin. The orange dashed line and red solid line represent three and four z-score thresholds, respectively. The y-axis shows the log2 ratio of ribosome protected fragments mapped downstream of the +1 frame stop codon over upstream (see Methods). The x-axis shows the total number of fragments aligned to the considered region of the PLEKHM2 mRNA.

**Figure S4 | Enhanced ARL8-independent localization of PLEKHM2-FS to cell tips. a.** ARL8-KO HeLa cells were transiently transfected with the indicated HA-tagged PLEKHM2 constructs for 48 h and analyzed for confocal immunofluorescence microscopy for the HA epitope (green), and endogenous LAMP1 (lysosomes, red). Nuclei were labeled with DAPI (blue). Cell edges were outlined by staining of actin with fluorescent phalloidin (not shown) and indicated by dashed lines. Scale bars: 20 μm. **b.** Quantification of the percentage of cells displaying localization of HA-PLEKHM2 variants to cell tips (left panel) or cytosol (right panel) from more than 150 cells per sample. Notice that HA-PLEKHM2-1-437 is less tip-localized and more cytosolic relative to HA-PLEKHM2-FS, suggesting a role for the transframe sequence in ARL8-independent localization of HA-PLEKHM2-FS to cell tips, indicative of interaction with kinesin-1.

**Figure S5 | Additional assays of PLEKHM2-KO hiPSC-CMs and WT hiPSC-CMs. a-b**. Quantification of average contraction and relaxation velocity (n = 10-12). **c-d**. qPCR of ANP and BNP transcripts, with fold ANP (NPPA) or BNP (NPPB) transcript calculated relative to WT (n = 12). **e.** Representative single trace of calcium transient. **f-h**. Quantification of amplitude, time to peak, and duration of calcium transients in hiPSC-CMs (n = 20-22). **i.** Western blot quantifications of p62 isoforms (n=4). **j.** Representative western blot showing p62 (H1 and H2 isoforms)^57^. All statistical tests performed were a one-way ANOVA (ns: p-value > 0.05, *: p-value < 0.05, **: p-value <0.01, ***: p-value <0.001, ****:p-value<0.0001).

## Methods

### Bioinformatic analysis

Reference sequences representing mammals, bird, amphibians and fish were used to generate sequences for each of these clades, in a manner previously reported^19^. For each clade, the sequences were translated to amino acids, aligned using MUSCLE^58^, and the amino acid alignments were used to guide codon-respecting nucleotide sequence alignments using EMBOSS tranalign^59^. For the MLOGD and synplot2 analyses, the alignments were mapped to the coordinates of a specific reference sequence by removing alignment columns that contained a gap character in the reference sequence. The reference sequences used were the coding regions of NCBI accession numbers NM_015164.4 (*Homo sapiens*), XM_417616.8 (*Gallus gallus*), XM_041569881.1 (*Xenopus laevis*) and NM_001130783.1 (*Danio rerio*) for the mammal, sauropsid, amphibian and teleost fish alignments, respectively. These alignments were analysed with synplot2^25^, using a 15-codon window size, and MLOGD^26^, using a 25-codon window size.

### Genome wide frameshifting analysis

Chordate mRNA sequences were downloaded from the NCBI RefSeq database in Nov 2017 and compiled into a BLAST nucleotide database^60^. For a given *Homo sapiens* gene name, the human RefSeq transcript with the longest protein-coding CDS out of those with an accession number beginning with "NM_" (i.e. manually curated mRNA sequences) was chosen as the reference sequence; if all accession numbers for the gene began with "XM_" (i.e. computationally annotated mRNA sequences) then one of these with the longest protein-coding CDS was used instead. Reference genes where the longest CDS was <100 codons or where the length of the annotated CDS was not a multiple of 3 were discarded. For each *Homo sapiens* reference mRNA sequence, the protein sequence was derived from the translated CDS, and tBLASTn (Altschul et al., 1990) was used to query the protein sequence against the aforementioned BLAST nucleotide database, using the parameter –qcov_hsp_perc 95 to ensure 95% coverage of the query sequence, and an e-value threshold of 0.001. Hits were filtered for ≥65% amino acid identity to the query sequence and, where multiple sequences from the same taxon were identified, only the hit with highest amino acid identity per taxon was kept. Reciprocal tBLASTn searches (with the same options as above) were performed using each hit from the first search as query sequence and the set of reference mRNAs as the subject sequences and only best reciprocal blast matches were retained. For each gene, the CDS sequences were extracted from each of the identified mRNA sequences, translated, and the amino acid sequences were aligned with MUSCLE^58^. Using tranalign from EMBOSS^59^, each amino acid alignment was used to guide a corresponding nucleotide alignment. Due to the 65% amino acid identity threshold, these alignments are often restricted to mammalian sequences but for highly conserved proteins they can also include sequences from other chordates. In total, alignments were generated for 17,922 human genes, contained a total of 1,586,159 sequences of which 75.1% were mammalian.

We identified all in-frame instances of YCC_UUU_CG in the 17,922 *Homo sapiens* CDSs, and then counted the number of sequences in the alignment, and the number containing a YCC_UUU_CG aligning exactly to the human YCC_UUU_CG. Out of 134 YCC_UUU_CG matches in human, only 7 were conserved in ≥95% of the aligned sequences. Five of these alignments were phylogenetically restricted (largely primate-specific), containing only 7–23 sequences. The remaining two were *ASXL1* (99 sequences, 98 with aligned YCC_UUU_CG) and *PLEKHM2* (114 sequences, 112 with aligned YCC_UUU_CG). Note that the procedure is not exhaustive: there may be instances of YCC_UUU_CG that are conserved in shorter genes (< 100 codons) or otherwise missed from the initial set of human reference sequences, conserved in some but not all of the sequences in an alignment (e.g. primate-specific), or conserved in presence but not in precise position across the MUSCLE sequence alignment, and such examples would have been missed by this analysis.

### Cell Culture and Transfections

For **Fig. 3**, HEK293T cells (ATCC) were maintained in DMEM supplemented with 10% FBS, 1 mM L-glutamine and antibiotics. For western blotting, cells were transfected with Lipofectamine 2000 reagent (Invitrogen) in 6-well plates using the 1-day protocol in which suspended cells are added directly to the DNA complexes in 6-well plates, with 1 mg of each indicated plasmid. The transfecting DNA complexes in each well were incubated with 5 × 10^5^ cells suspended in 3 ml DMEM + 10% FBS and incubated for 36 h at 37°C in 5% CO_2_.

For GFP-Trap transfections, 1 x 10^7^ cells were forward-transfected with Lipofectamine 2000 reagent in 15 cm Petri dishes with 10 mg of each indicated plasmid. Cells were incubated for 48 h at 37°C in 5% CO_2_.

For dual luciferase assays, cells were transfected with Lipofectamine 2000 reagent, again using the 1-day protocol described above. The following were added to each well: 25 ng of each plasmid plus 0.2 μl Lipofectamine 2000 in 25 μl Opti-Mem (Gibco). The transfecting DNA complexes in each well were incubated with 3 × 10^4^ cells suspended in 50 μl DMEM + 10% FBS at 37°C in 5% CO_2_ for 20 h.

For polyamine experiments, cells were transfected using Lipofectamine 2000 reagent as described above. Suspended cells were supplemented to a final concentration of 1 mM aminoguanidine hydrochloride (Sigma), or 1 mM aminoguanidine hydrochloride plus 2 mM spermidine (Sigma) before adding to transfecting DNA complexes. Luciferase activities were measured 20 h after spermidine treatment.

For **Fig. 4**, HeLa and HEK293T cells were cultured in DMEM (Quality Biological, #112-319-101), supplemented with 2 mM L-glutamine (GIBCO, #25030081), 10% fetal bovine serum (BSA) (Corning, # 35-011-CV), 100 U/ml penicillin-streptomycin (GIBCO, # 15140122) (complete DMEM) in a 37°C incubator (5% CO2, 95% air). HeLa cells grown on 6-well plates were transiently transfected with 2 μg plasmid DNA using 8 μl Lipofectamine 3000 (Invitrogen, #L3000001), according to the manufacturer’s instructions. Approximately 24 h after transfection, 40,000 cells were replated onto 12-mm coverslips coated with collagen (BioTechne, #3442-050-01). The remaining unseeded cells were used for SDS-PAGE and immunoblotting. Cells were then cultured for an additional 24 h before fixation and immunofluorescent labeling. For co-immunoprecipitation experiments, HEK293T cells grown on 6-well plates were transiently transfected for 48 h with 1.5 μg of each plasmid DNA and 8 μl Lipofectamine 3000 (Invitrogen, #L3000001) according to the manufacturer’s instructions.

### Expression constructs

*PLEKHM2* fused dual luciferase constructs (for immunoblotting) were generated by either 1-step or 2-step PCR on *PLEKHM2* gBlock1 (IDT) using primer sequences which incorporated 5′ *Xho*I and 3′ *Bgl*II restriction sites (outlined in Supplementary Table 1). PCR amplicons were digested with *Xho*I / *Bgl*II and cloned into *Psp*XI / *Bgl*II digested pDlucV2.0. pDlucV2.0 is a version of pDluc^31^ generated by introducing silent mutations into the *Renilla* coding sequence to disrupt two potential donor splice sites (TGGgtaagt).

PLEKHM2-X-GFP and PLEKHM2-K-X-GFP were synthesized as gBlocks with flanking 5′ *Sac*I and 3′ *Bam*HI restriction sites and cloned into pcDNA3.4. *PLEKHM2* unfused dual luciferase constructs (for dual luciferase assay) were generated by either 1-step or 2-step PCR on a *PLEKHM2* G Block (IDT) using primer sequences which incorporated 5′ *Xho*I and 3′ *Bgl*II restriction sites (outlined in Supplementary Table 1). PCR amplicons were digested with *Xho*I / *Bgl*II and cloned into *Psp*XI / *Bgl*II digested pSGDlucV3.0 (Addgene 119760)^33^. *OAZ1* dual luciferase expression constructs were described previously^61^. All clones were verified by Sanger sequencing (Eurofins). All constructs used for cell biology assays were directly synthesized by Twist Biosciences for overexpression in mammalian cells.

### Antibodies and chemicals

The following primary antibodies were used for immunoblotting (IB) and/or immunofluorescence microscopy (IF) (catalog numbers, sources and working dilutions are in parentheses): rat anti-HA (12158167001, Roche, IF 1:500, IB 1:1,000), mouse anti-LAMP1 (H4A3, DSHB, IF 1:500), rabbit anti-LAMTOR4 (13140, Cell Signaling, IF 1:500), rabbit anti-myc-tag (2272, Cell Signaling, IB 1:1,000), mouse anti-tubulin-HRP (sc-32293, Santa Cruz, IF 1:1,000). Secondary antibodies: HRP-conjugated goat anti-rat IgG (H+L) (112-035-003, Jackson Immuno Research, IB 1:5,000), HRP-conjugated goat anti-rabbit IgG (H+L) (111-035-003, Jackson Immuno Research, IB 1:5,000), Alexa Fluor 555-conjugated donkey anti-mouse IgG (A-31570, Thermo Scientific, IF 1:1,000), Alexa Fluor 555-conjugated goat anti-rabbit IgG (A-21428, Thermo Scientific, IF 1:1,000), Alexa Fluor 488-conjugated goat anti-rat IgG (A-11006, Thermo Scientific, IF 1:1,000). Cell edges were outlined by staining of actin with Alexa FluorTM 647-conjugated phalloidin (A22287, Thermo Scientific, IF 1:500).

### Dual Luciferase Assay

Relative light units were measured on a Veritas Microplate Luminometer with two injectors (Turner Biosystems). Transfected cells were lysed in 15 μl of 1 × passive lysis buffer (PLB: Promega) and light emission was measured following injection of 50 μl of either *Renilla* or firefly luciferase substrate^62^. Recoding efficiencies were determined by calculating relative luciferase activities (firefly/*Renilla*) from test constructs and dividing by relative luciferase activities from replicate wells of in-frame-control constructs. Four replicate biological samples were assayed each with four technical repeats. Statistical significance was determined using a two-tailed, homoscedastic Student’s *t*-test.

### SDS-PAGE and immunoblotting

For **Fig. 3**, transfected cells were lysed in 100 ml 1 × PLB. Proteins were resolved by SDS-PAGE and transferred to nitrocellulose membranes (Protran), which were incubated at 4°C overnight with primary antibodies. Immunoreactive bands were detected on membranes after incubation with appropriate fluorescently labeled secondary antibodies using a LI-COR Odyssey® Infrared Imaging Scanner.

For **Fig. 4**, cells were washed twice with ice-cold phosphate-buffered saline (PBS; Corning, #21-040-CM), scraped from the plates in 1x Laemmli sample buffer (1x LSB) (Bio-Rad, #1610747) supplemented with 2.5% v/v 2-mercaptoethanol (Sigma-Aldrich, #60-24-2), heated at 95°C for 5 min, and resolved by SDS-PAGE. Gels were blotted onto nitrocellulose membrane and blocked with 5% w/v non-fat milk in Tris-buffered saline, 0.1% v/v Tween 20 (TBS-T) for 20 min. Membranes were sequentially incubated with primary antibody and secondary HRP-conjugated antibody diluted in TBS-T. SuperSignal West Dura Reagents (Thermo Fisher, #34075) were used for detection of the antibody signal with a Bio-Rad Chemidoc MP imaging system. β-tubulin was used as a control for sample loading.

For **S5i-j**, equal amounts of protein were loaded (normalized by a BCA assay) and run on a SDS-PAGE gel. Protein was then transferred to a PVDF membrane that was blocked with 5% skim milk for 1 hour at 37°C before an overnight incubation with primary antibodies at 4°C. The next day, secondary antibodies were incubated at room temperature for 1 hour prior to ECL chemiluminescence measurement.Signal intensity analysis was performed in ImageJ.

### Immunoprecipitation

For **Fig. 3**, Cells were lysed in 700 ml PLB and then incubated with 20 ml of protein G Agarose beads plus anti-HA (3 mg) overnight at 4°C with gentle rocking. The beads were washed with ice-cold 1 × PLB buffer and then removed from the beads by boiling for 5 min in 2 × SDS-PAGE sample buffer for SDS-PAGE and western blotting. For GFP immunoprecipitation, GFP-Trap (Chromtek) was used following the manufacturer’s instructions. Briefly, cells were lysed in 100 ml NP-40 lysis buffer (10 mM Tris/Cl pH 7.5; 150 mM NaCl; 0.5 mM EDTA; 0.5% NP-40) then 95 ml lysate was diluted to 700 ml in dilution buffer (10 mM Tris/Cl pH 7.5; 150 mM NaCl; 0.5 mM EDTA) before incubation with 20 ml of GFP-Trap beads for 1 h at 4°C with gentle rocking. The beads were washed with ice-cold dilution buffer and then removed from the beads by boiling for 5 min in 2 × SDS-PAGE sample buffer for SDS-PAGE and Coomassie staining.

For **Fig. 4**, transfected HEK293T cells were lifted in ice-cold PBS and centrifuged at 500 *g* for 5 min. The pellet was washed twice with ice-cold PBS and lysed in 10 mM Tris-Cl pH 7.5, 150 mM NaCl, 0.5 mM EDTA, 0.5% v/v Nonidet P40 supplemented with a protease inhibitor cocktail (Roche). Cell lysates were clarified by centrifugation at 17,000 *g* for 10 min and the supernatant (10% was saved as input) was incubated with anti-myc magnetic agarose beads (Thermo Fisher, #88842) at 4°C for 2 h. After three washes with 10 mM Tris-Cl pH 7.5, 150 mM NaCl, 0.5 mM EDTA, the immunoprecipitates were eluted with 1x LSB at 95°C for 5 min. The immunoprecipitated samples and inputs were analyzed by SDS-PAGE and immunoblotting.

### Immunofluorescence microscopy

Cells were seeded on collagen-coated coverslips in 24-well plates at 40,000 cells per well in regular culture medium. After 24 h, cells were fixed in 4% w/v paraformaldehyde (Electron Microscopy Sciences, #15700) in PBS for 20 min, permeabilized and blocked with 0.1% w/v saponin, 1% w/v BSA (Gold Bio, #A-421-10) in PBS for 20 min, and sequentially incubated with primary and secondary antibodies diluted in 0.1% w/v saponin, 1% w/v BSA in PBS for 30 min at 37°C. Coverslips were washed three times in PBS and mounted on glass slides using Fluoromount-G (Electron Microscopy Sciences, #17984-24) with DAPI. Z-stack cell images were acquired on a Zeiss LSM 900 inverted confocal microscope (Carl Zeiss) using a Plan-Apochromat 63X objective (NA = 1.4) with or without Airyscan detection. Maximum intensity projections were generated with Zeiss ZEN Black software, and final composite images were created using ImageJ/Fiji (https://fiji.sc/).

### Ribosome profiling

Gencode v.46 transcript ENST00000375799 (4134 bp) was used as a representative mRNA of PLEKHM2. The frameshift cassette (UCC_UUU_CGG) is located between the coordinates 1545 -1553, and frameshifted ribosomes terminate at an early stop codon in the new reading frame (+1) at coordinates 1789 -1791 (*TAG*). Intuitively, PRF is expected to reduce the number of ribosome protected fragments (RPFs) aligning downstream of the frameshift-introduced stop codon, and as such the reduction in read density reflects the efficiency of PRF.This relationship can be mathematically represented as the ratio between read density downstream over the one upstream the frameshift cassette.

To limit the effect of aberrant pauses around translation initiation and termination sites, the 100 nt downstream and upstream, respectively, have been discarded. In a similar reasoning, the region from 50 nt upstream the frameshift cassette and up to 50 nt downstream the frameshift-introduced stop codon has been excluded from the analysis. Specifically, the region between coordinates 340 -1494 was used to estimate the density of RFPs upstream the frameshift cassette, while coordinates 1842 -3199 identify the region used to estimate the density downstream of the frameshift-introduced stop codon. TPM (transcripts per kilobase million) values for corresponding regions were obtained from RiboCrypt (https://ribocrypt.org/) for each individual dataset, and to calculate the ratio the mean value of TPM for each region was considered. The ratios were then log2-transformed for the further steps of the analysis.

The classification by tissue of origin was based on the curated metadata available in RiboSeq Data Portal (https://rdp.ucc.ie). To assess the statistical significance of the ratio variation, we used a sliding window Z-score approach similar to what was described earlier^63^. Briefly, the log2 ratios from each study were sorted by ascending values of “total counts” -that is to say, the sum of all the raw counts mapping to each nucleotide position in the regions before and after the frameshift cassette. Then the ratios were grouped into bins of size 300 and descriptive statistics (mean and standard deviation) were calculated and stored. The window was then shifted with a step of 10 and the statistics recomputed, until the end of the available ratios. At that point, each ratio had a collection of means and standard deviations which were averaged and used to obtain the Z-score value for that specific log2 ratio.

To represent thresholds of Z-scores of 3 and 4 over the log_2 ratio distributions, we used the binning-and-sliding window approach mentioned before to calculate the standard deviations and means for each value, but with bin size of 50 and step of 10. The intervals between 3 and 4 standard deviations from the mean were then computed for each datapoint and plotted on the graph as two threshold lines (Fig. S3).

### iPSC cell culture, differentiation, purification and inoculation

iPSCs were generated from the ZZUNEUi022-A cell line^64^ which was established from male urine cells. PLEKHM2-KO hiSPECs were generated with a single guide RNA (sgRNA) targeting the second exon of *PLEKHM2* and verified by sequencing and western blot^48^. Matrigel coated 6-well plates were seeded with iPSCs at 10,000 cells per well and cultured in PSCeasy medium at 5% carbon dioxide at 37°C. hiSPCs were differentiated into cardiomyocytes (hiPSC-CMs) using a small molecule-based approach described previously^49^. On the 30th day of differentiation, lentivirus is added to the hiPSC-CMs at an MOI of 4. After a week post-infection, cells were assayed.

### Contractility measurements

Dissociated cardiomyocytes were reseed onto micropatterned hydrogels with stiffness of 10 kPa. These single cell patterns were arranged in a 1 mm by 1 mm grid, spaced to minimize the risk of mechanical interaction between adjacent cells. Single cell contraction force was assessed from single cells using Zeiss fluorescence microscope . We acquired two types of videos via microscopy. Videos of cardiomyocyte contractions were then collected in fluorescence channels, capturing cardiac contractions at 30 frames per second with a 40x objective for 10 seconds. Cells were imaged in bright field to determine individual cells and beating rates. Video of the movement of fluorescent beads in the substrate caused by the contraction of cardiomyocytes through the fluorescence channel. Raw video was extracted using Hcell^65^ software. Briefly, the contraction force was calculated from Video of the movement of fluorescent beads in the substrate and we calculated the magnitude of force vectors (F) from traction stresses. The velocity and contractile force corresponding to each time point are calculated from each frame of the two videos, and the corresponding velocity and contractile force curves are drawn based on these data.

### qPCR of ANP and BNP

mRNA was isolated from the hiSPC-CMs using TRIzol and reverse-transcribed into cDNA with PrimeScript RT Master Mix. Quantitative real-time PCR was performed on samples using a QuantStudio 3 instrument along with TB Green Premix Ex Taq II PCR mix. GAPDH was used as a housekeeping gene to calculate the relative gene expression using the double delta CT method. The primer sequences used for GAPDH were: GGAGCGAGATCCCTCCAAAAT and GGCTGTTGTCATACTTCTCATGG, ANP (NPPA): ACAATGCCGTGTCCAACGCAGA and CTTCATTCGGCTCACTGAGCAC, and BNP (NPPB): TCTGGCTGCTTTGGGAGGAAGA and CCTTGTGGAATCAGAAGCAGGTG

### Calcium transient measurements

Calcium handling was measured as previously described^48^. Briefly, the green fluorescent calcium-modulated protein 6 fast type (GCaMP6f) calcium imaging system was used to track the calcium transients in hiPSC-CMs. Using the Plexithermo system, spontaneous calcium transients of individual cells on an OLYMPUS IX73 microscope at 37°C were measured and the data were analyzed using the ImageJ software.

## Acknowledgements

Y.A.K. was supported by a Knight-Hennessy Scholarship, a NSF Graduate Research Fellowship Program award, and an NIH F31 grant (F31MH134477). A.E.F. and J.W. were supported by Wellcome Trust Senior Research Fellowships (106207/Z/14/Z, 220814/Z/20/Z) and a European Research Council grant (646891) to A.E.F. J.F.A. was supported by an Irish Research Council Advanced Laureate Award (IRCLA/2019/74). P.V.B. was supported by Science Foundation Ireland Frontiers for the Future Award (20/FFP-A/8929) and SFI-HRB-Wellcome Trust Biomedical Research Partnership Investigator Award in Science (210692/Z/18/Z). G.C. is supported by Science Foundation Ireland Centre for Research Training in Genomics Data Science [18/CRT/6214]. Work in the laboratory of X.L. was supported by the National Natural Science Foundation of China. Work in the J.S.B. lab was supported by the Intramural Program of NICHD, NIH (project ZIA HD001607). J.M.M. and I.J. were supported by the National Human Genome Research Institute (NHGRI) of the U.S. National Institutes of Health (NIH) under award number (U24HG007234). The content is solely the responsibility of the authors and does not necessarily represent the official views of the National Institutes of Health. J.M.M. was also supported by the Wellcome Trust (108749/Z/15/Z) and the European Molecular Biology Laboratory (EMBL). Ensembl is a registered trademark of EMBL. For the purpose of open access, the authors have applied a CC BY public copyright license to any Author Accepted article version arising from this submission. The authors would like to acknowledge Michael Z. Palo for discussions related to this manuscript.

## Notes

### Competing Interest Statement

The authors have declared no competing interest.

### Summary of Updates

This is a new and updated version of this manuscript that has been resubmitted to Nature. This new version contains various new phenotypic data related to the frameshift element's role in myocardial contraction

