## Supplementary figures and images for "Programmed ribosomal frameshifting during *PLEKHM2* mRNA decoding generates a constitutively active proteoform that supports myocardial function"

### S1

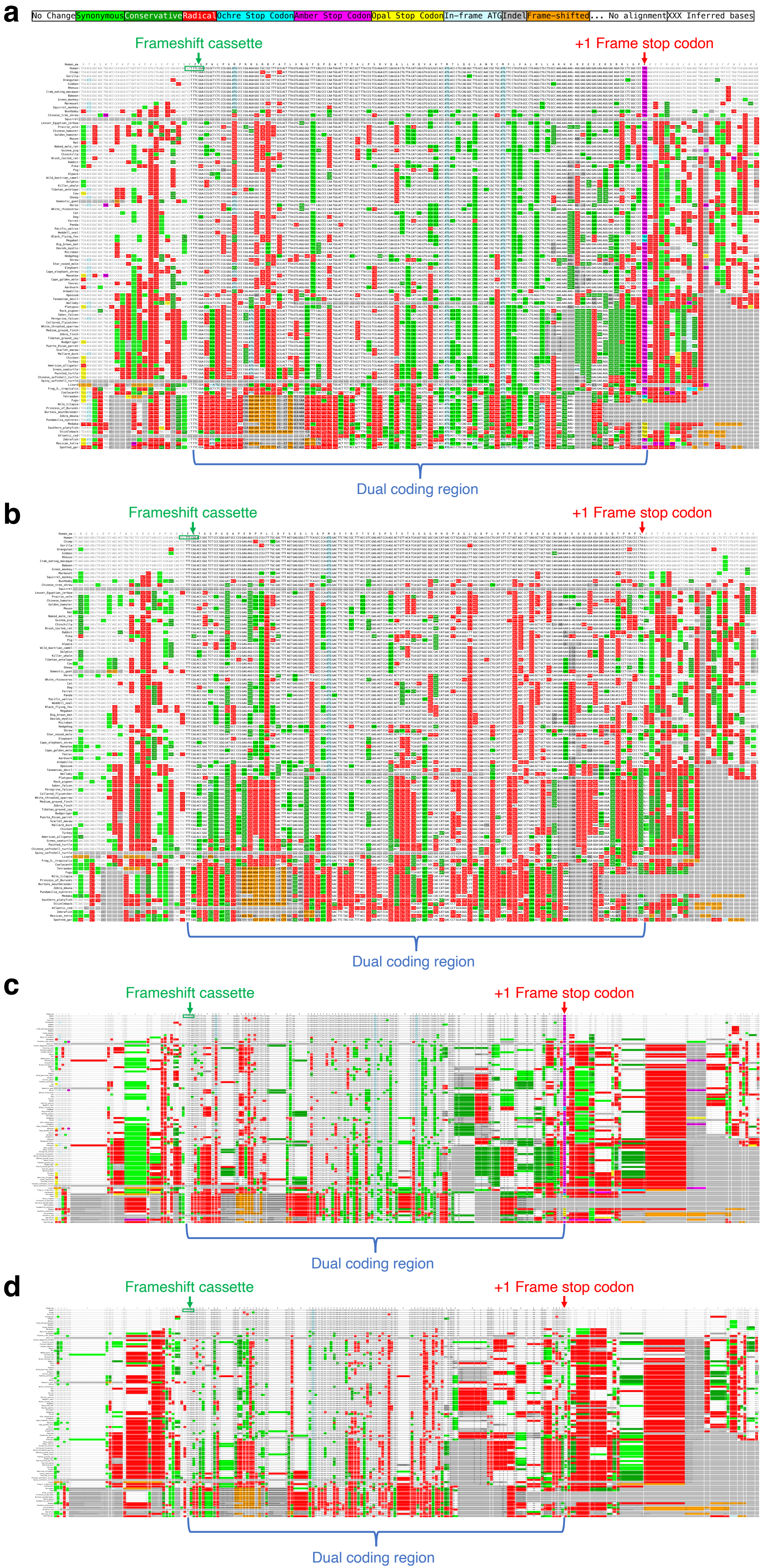

### S2

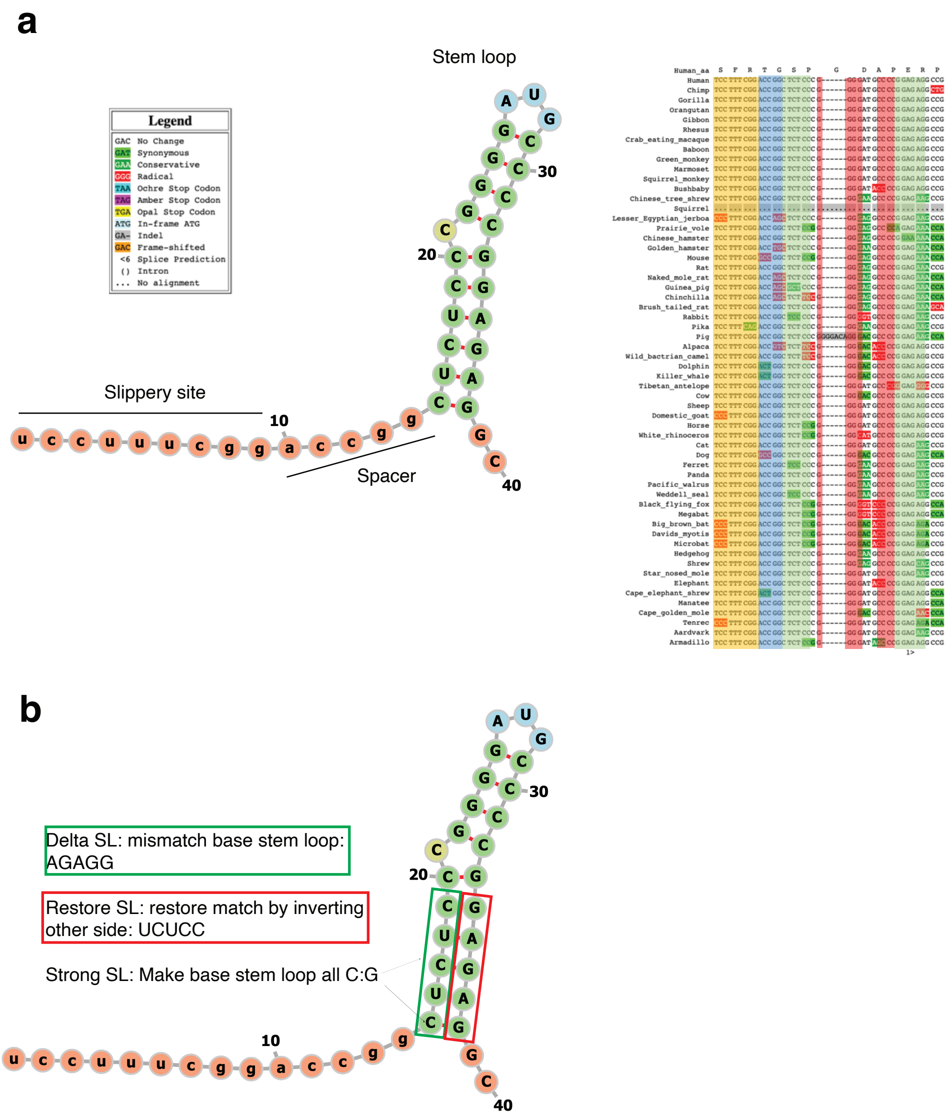

### S3

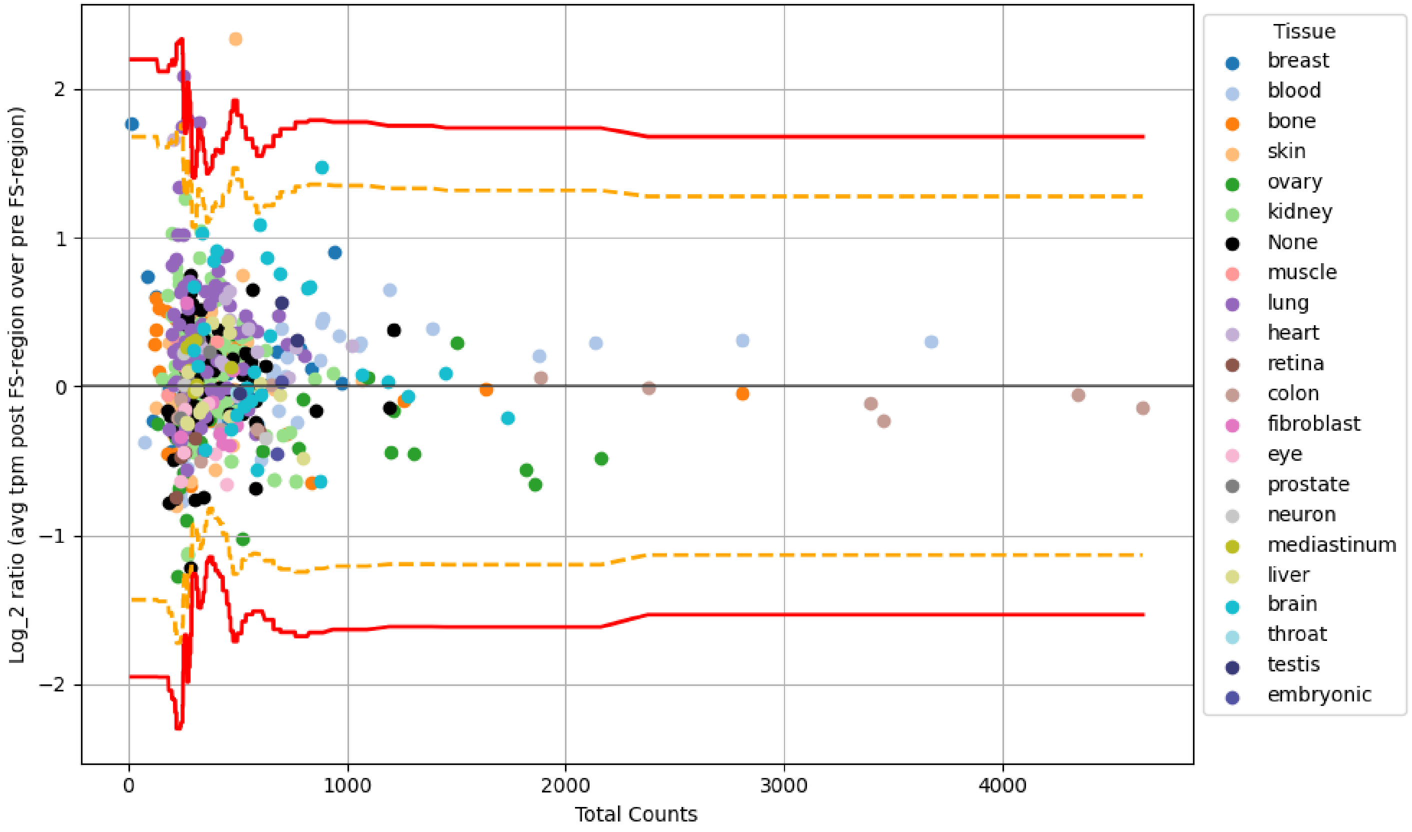

### S4

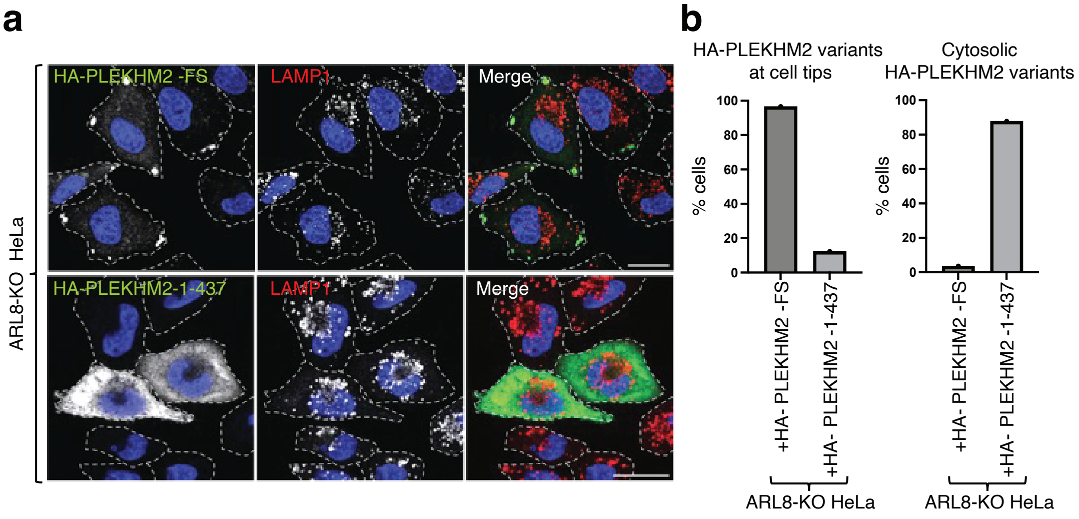

### S5

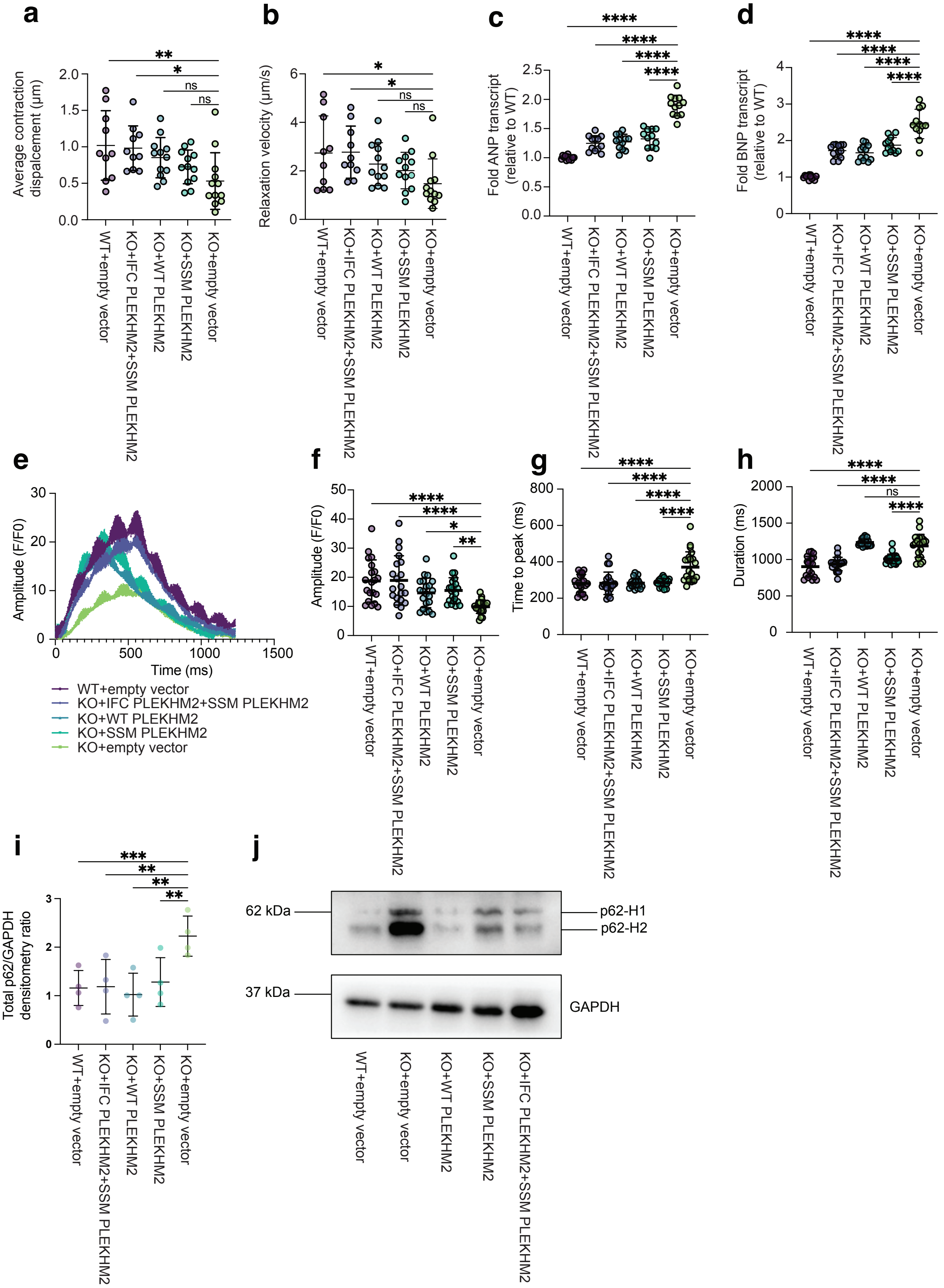
